# KDM6A/UTX promotes spermatogenic gene expression across generations but is dispensable for male fertility

**DOI:** 10.1101/2022.10.27.513976

**Authors:** Benjamin William Walters, Shannon R Rainsford, Nicolas Dias, Xiaofang Huang, Dirk G de Rooij, Bluma J Lesch

## Abstract

Paternal chromatin undergoes extensive structural and epigenetic changes during mammalian spermatogenesis, producing sperm that contain an epigenome optimal for the transition to embryogenesis. Histone modifiers play an important role in this process by encoding specialized regulatory information in the sperm epigenome. Lysine demethylase 6a (KDM6A) promotes gene activation via demethylation of H3K27me3, a developmentally important repressive modification abundant throughout the epigenome of sperm and embryonic stem cells. Despite its developmental importance in pluripotent cells and germ cell progenitors, the function of KDM6A during spermatogenesis has not been described. Here, we show that *Kdm6a* is transiently expressed in the male germline in late spermatogonia and during the early stages of meiotic entry. Deletion of *Kdm6a* in the male mouse germline (*Kdm6a* cKO) yielded a modest increase in sperm head defects but did not affect fertility or the overall progression of spermatogenesis. However, hundreds of genes were deregulated upon loss of *Kdm6a* in spermatogenic cells and in an immortalized spermatogonia cell line (GC-1 spg) with a strong bias towards downregulation. Single cell RNA-seq revealed that most of these genes were deregulated in spermatogenic cells at the same stage when *Kdm6a* is expressed and encode epigenetic factors involved in chromatin organization and modification. A subset of these genes was persistently deregulated in the male germ line across two generations of offspring of *Kdm6a* cKO males. Our findings highlight KDM6A as a transcriptional activator in the mammalian male germline that is dispensable for spermatogenesis but important for safeguarding gene regulatory state intergenerationally.

**Author summary:** Offspring viability and fitness relies upon the development of functional sperm and the integrity of information that they carry. Chromatin is modified and remodeled extensively throughout spermatogenesis to facilitate meiosis, DNA compaction, and to encode gene regulatory information for the next generation. In mice, a paternal germline lacking KDM6A, a histone modifier, yields offspring with reduced lifespans and increased cancer risk. How KDM6A functions in the paternal germline to support offspring health is unknown. Here, we show that *Kdm6a* expression is limited to a distinct developmental interval when differentiated spermatogonia transition from mitosis to meiosis. During this timepoint, KDM6A acts as a transcriptional activator for hundreds of genes, many of which encode meiotic factors and epigenetic modifiers. Nevertheless, this activity is dispensable for overall spermatogenesis and fertility. Surprisingly, we find a significant overlap in germline transcriptomes of *Kdm6a* cKO mice and wildtype offspring. We propose that KDM6A encodes gene regulatory information in the male germline that is retained across generations.

## Introduction

Chromatin states are a major determinant of gene expression. Posttranslational histone modifications play a central role in defining and regulating chromatin state, and histone modifying enzymes are critical factors in guiding gene expression programs. The importance of histone modification dynamics is particularly evident in spermatogenesis, when chromatin is dramatically remodeled and modified to facilitate precise temporal gene expression programs and to generate a viable epigenome for the next generation. One of the best characterized histone modifications in the male germline is trimethylation of histone H3 at lysine 27 (H3K27me3), which is catalyzed by the Polycomb repressive complex 2 (PRC2) (1, 2). Extensive studies in multiple cell types point to a role for PRC2 and H3K27me3 in limiting promiscuous gene expression. Notably, H3K27me3 is enriched at genes encoding developmental regulators in mouse embryonic stem cells and has been shown to support ground-state pluripotency (3, 4). In male germ cells, loss of PRC2 leads to derepression of somatic and stage-specific meiosis genes and ultimately causes sterility (5). H3K27me3 is also one of the two histone modifications that define bivalency, a unique chromatin state specified by co-occupancy of H3K27me3 and the activation mark H3K4me3 and thought to poise repressed germline genes for later activation in the developing embryo. Several studies have linked paternally-derived H3K27me3 epimutations with detrimental developmental consequences in offspring (6-9). Therefore, modulation of the levels and distribution of H3K27me3 in the male germline is critical for fertility and for facilitating appropriate offspring development.

The X-linked gene *Kdm6a* (also known as *Utx*) counterbalances PRC2 activity by demethylating H3K27me2/3 to derepress genes and promote transcription (10). KDM6A was initially described as a critical developmental factor in *C. elegans* and zebrafish through its regulation of Hox gene activity (11, 12). KDM6A has since been shown to play important roles in diverse cellular processes, including embryonic stem cell differentiation, aging, cellular reprogramming and cellular development (13-22). KDM6A also has non-catalytic activity in regulation of enhancer activation via association with the MLL3/4 complex (37). In aggregate, existing data highlight KDM6A as an important transcriptional regulator with complex roles in the establishment of gene expression programs.

In addition to its role in development, missense and truncating mutations in *KDM6A* have been identified across a broad range of human cancers, suggestive of a function for KDM6A as a tumor suppressor (23). Loss of *Kdm6a* in the mouse paternal germline potentiates cancer risk in genetically wildtype offspring, implying that some of the tumor suppressive epigenetic effects of KDM6A are heritable (8). This phenotype suggests a model in which KDM6A primes the sperm epigenome with gene regulatory information that limits malignant transformation. How KDM6A functions across spermatogenesis in the parental germline to confer these effects in the next generation is not fully understood.

In this study, we investigate the function of KDM6A in the male germline, with a particular focus to understanding its effects on gene regulation during spermatogenesis. We show that *Kdm6a* expression is limited to late spermatogonia and the early stages of meiotic prophase, in contrast to the broad expression of most histone demethylase genes during spermatogenesis. KDM6A acts as a transcriptional activator predominantly during these early stages of meiosis, where it targets genes encoding chromatin remodelers and regulators of chromosome organization, among others. KDM6A mildly impacts chromatin accessibility and does not alter global H3K27me3 levels, suggesting that KDM6A may act primarily via alternative mechanisms in the male germline. Despite its effects on gene expression, loss of *Kdm6a* did not overtly impact the progression of spermatogenesis, although it did cause a mild increase in sperm head defects. Intriguingly, we identified a subset of genes positively regulated by KDM6A that were persistently deregulated in the testes of wildtype offspring of *Kdm6a* cKO males. Overall, our data support a model in which KDM6A resets gene regulatory information in the male germline that is not essential for fertility but can affect gene expression across generations.

## Results

### KDM6A is transiently enriched in primordial germ cells and in spermatogenic cells at meiotic entry

Although *Kdm6a* is reported to be ubiquitously expressed in mice, including in reproductive organs (24), its expression pattern across gonad development and adult spermatogenesis has not been carefully defined. To broadly address this question, we first measured the abundance of *Kdm6a* transcripts using bulk RNA sequencing at three time points in post-migratory primordial germ cells (PGCs) (25), and in enriched populations of meiotic (pachytene spermatocyte) and post-meiotic (round spermatid) cells, as well as whole testes. Expression of *Kdm6a* was detectable in all samples but was most enriched in PGCs relative to adult male germ cells (**Fig 1A**). The enrichment of *Kdm6a* transcript in PGCs is consistent with a previous report implicating KDM6A as a regulator of epigenetic reprogramming in PGCs (17). *Kdm6a* transcript levels were higher in whole adult testis compared to either pachytene spermatocytes or round spermatids. A similar trend was evident at the protein level, where Western blotting revealed moderate KDM6A protein expression in whole testis that was reduced in post-meiotic cells and almost undetectable in meiotic cells (**Fig 1B**), suggesting that a population of testicular somatic cells or pre-pachytene germ cells expresses *Kdm6a* at a level higher than either pachytene spermatocytes or round spermatids. Supporting this hypothesis, *Kdm6a* transcript levels were significantly higher in an immortalized spermatogonia-derived cell line (GC-1 spg, hereafter called GC1s) (26) relative to PGCs, whole adult testis, pachytene spermatocytes and round spermatids (**Fig 1A**).

**Fig 1.**
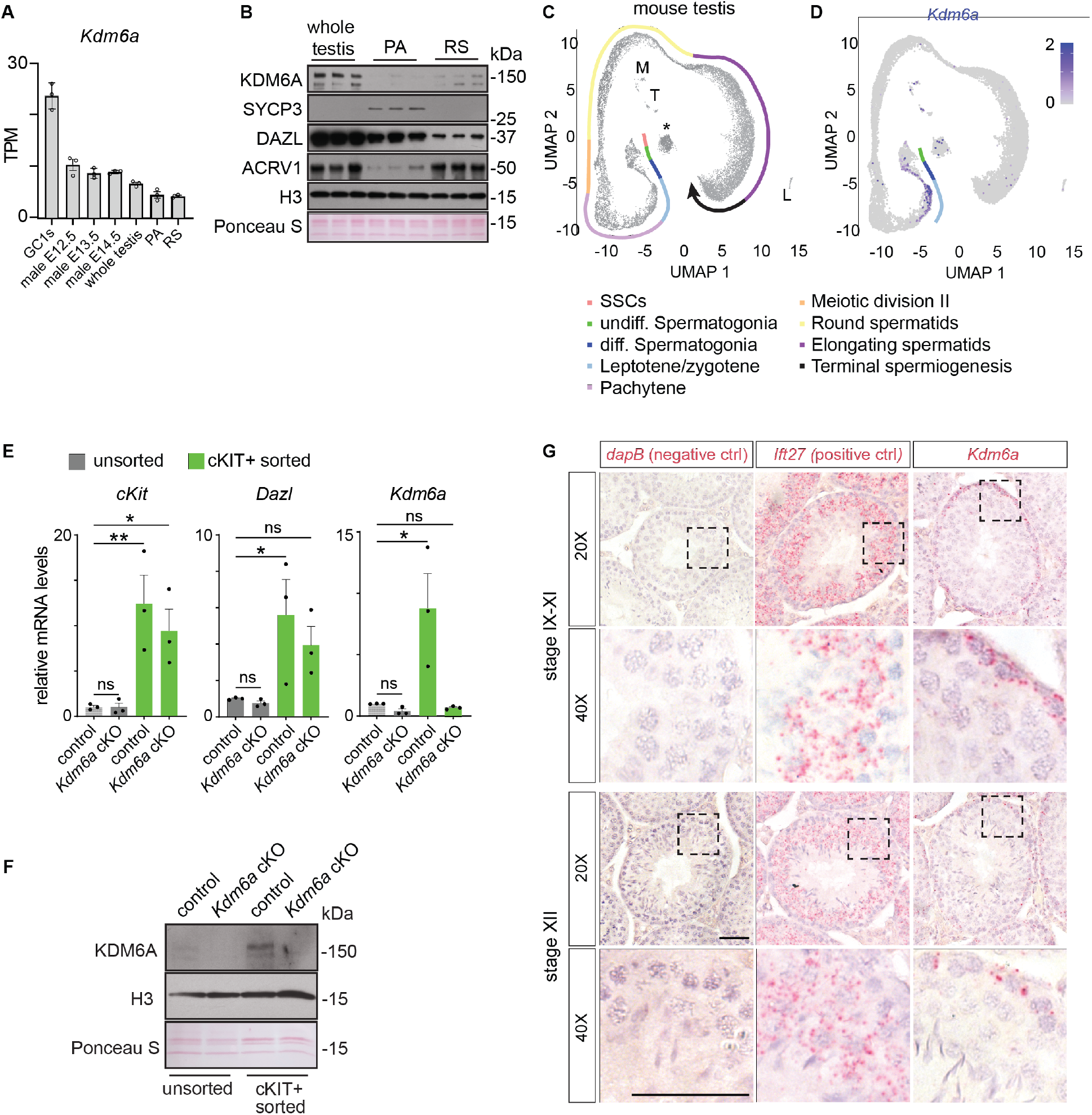
Expression analysis of KDM6A in the male germline. (**A**) Transcripts per million (TPM) for *Kdm6a* in the GC1-SPG cell line, primordial germ cells, whole adult testis, pachytene-enriched (PA) and round spermatid-enriched male germ cells (RS). (**B**) Western blotting in male germ cells for KDM6A and markers of meiotic (SYCP3) and post-meiotic (ACRV1) cells. DAZL is as a broad male germ cell marker and histone H3 is as a loading control. (**C**) UMAP of single cell RNA-seq (scRNA-seq) data from whole mouse testis. Distinct cell populations are indicated by the colored line. M = macrophages, T = telocytes, L = Leydig cells, * = unassigned. (**D**) UMAP showing the expression pattern of *Kdm6a* across testis cell populations detected by scRNA-seq. (**E**) RT-qPCR in unsorted and cKIT-sorted cells derived from testes of control and *Kdm6a* cKO mice. ns = not significant, * = P ≤ 0.05, ** = P ≤ 0.01. (**F**) Western blotting for KDM6A in unsorted and cKIT-sorted cells derived from testes of control and *Kdm6a* cKO mice. (**G**) Brightfield micrographs showing in situ hybridization for the indicated transcript (pink) in tissue sections of seminiferous tubules co-stained with hematoxylin. Dashed boxes indicate the regions captured at high magnification below. Scale bar = 50μm.

To more precisely identify the cell type with strongest *Kdm6a* expression in adult testes, we generated a single cell RNA-seq (scRNA-seq) dataset from adult mouse testis and examined the dynamics of *Kdm6a* gene expression across spermatogenic development. Graph-based clustering using Seurat (27) defined seventeen distinct clusters representing the known trajectory of spermatogenic maturation along with most testicular somatic cell types, with the exception of Sertoli cells (**Fig 1C, S1A and S1B**). Consistent with our bulk RNA and protein data, *Kdm6a* expression was not uniform across testicular cell types. *Kdm6a* was highly expressed in several somatic cell populations, including macrophages, Leydig cells and telocytes (**Fig 1D**). Among germ cells, *Kdm6a* mRNA expression was almost exclusive to a defined developmental interval encompassing late spermatogonia and the early stages of meiotic prophase. *Kdm6a* transcripts were most abundant in intermediate/spermatogonia B, differentiated spermatogonia, and leptotene/zygotene spermatocytes. We confirmed this expression profile with publicly available scRNA-seq datasets from mouse (28) and human (29) (**Fig S1C and S1D**). To validate this expression pattern, we enriched for late spermatogonia/early meiotic prophase germ cells from whole testes by flow cytometry for cKIT+ cells (**Fig S1D**) (30) and performed RT-qPCR and Western blotting for *Kdm6a*. We confirmed a strong enrichment for *Kdm6a* mRNA and protein relative to unsorted cells and cKIT+ cells isolated from testes of a *Kdm6a* germline conditional knockout (cKO, see next section) (**Fig 1E and 1F**). Further confirming this expression pattern, in situ hybridization revealed an enrichment for *Kdm6a* transcript in the basal compartment of the seminiferous tubule where spermatogonia and early meiotic cell populations reside (**Fig 1G and Fig S1E**). This was particularly evident in tubules at stage IX-XII of the spermatogenic cycle.

*Kdm6a* has two homologs in the mouse genome, *Uty* and *Kdm6b*. We queried our scRNA-seq dataset to see if *Uty* and *Kdm6b* had similar gene expression patterns to *Kdm6a. Uty* was weakly expressed in undifferentiated spermatogonia and early meiotic cells while *Kdm6b* expression was generally enriched in spermatogonia, implying that both homologs are expressed in overlapping but not identical cell populations compared to *Kdm6a* in the adult mouse testis (**Fig S1F**). To determine if the expression pattern of *Kdm6a* is unique among histone demethylase genes, we also assessed the expression patterns of eighteen other lysine demethylases using our scRNA-seq data. Seven lysine demethylases exhibited broad expression patterns across spermatogenesis, while eleven exhibited stage-specific expression profiles that did not fully overlap with *Kdm6a* (**Fig S1D**). The expression pattern of *Kdm7b* was most similar to *Kdm6a* during early spermatogenesis but was also detectable in round spermatids. Interestingly, *Kdm7b* can demethylate mono- and di-methylated, but not tri-methylated H3K27 (31), implying that *Kdm6a* and *Kdm7b* may have complementary, non-redundant functions during spermatogenesis. Overall, *Kdm6a* has a unique and transient expression pattern during spermatogenesis.

Together, these data demonstrate that contrary to previous assumptions, *Kdm6a* is not ubiquitously expressed in spermatogenesis. KDM6A is temporally regulated during spermatogenesis and is most strongly expressed from late stages of spermatogonial differentiation to the earliest stages of meiotic prophase, indicating a potential function for KDM6A during meiotic entry.

### *Kdm6a* is dispensable for spermatogenesis

To evaluate the function of KDM6A in spermatogenesis, we generated mice lacking *Kdm6a* specifically in germ cells. These mice carry a conditional allele of *Kdm6a* (22) along with the germline-specific *Ddx4*-Cre (32), resulting in deletion of the third *Kdm6a* exon and yielding a truncated non-functional protein specifically in the germline. In *Kdm6a* cKO males, *Kdm6a* transcript and protein expression was significantly decreased in cKIT+ cells and whole testis lysates compared to control, confirming the efficacy of the knockout (**Fig 1E, 1F and S2A**). There was no change in the transcript levels of *Uty, Kdm6b*, or *Kdm7b*, in *Kdm6a* cKO testes, indicating that KDM6A homologs are not compensating for *Kdm6a* loss at the transcript level (**Fig S2B**).

We next asked if *Kdm6a* loss in the male germ line caused any phenotypic abnormalities. Males from our *Kdm6a* cKO mouse line were previously shown to be normally fertile based on number of offspring (8). We found that *Kdm6a* cKO testes were comparable in weight and size to control mice when normalized to body weight (**Fig 2A and 2B**). Histologically, all major spermatogenic cell types were present and morphologically normal in *Kdm6a* cKO seminiferous tubules (**Fig 2C**). Overall protein levels and spatiotemporal expression of markers for spermatogonia (LIN28, SALL4, STRA8 and CHD4), spermatocytes (SYCP3) and general spermatogenic cells (DAZL) were also comparable between control and *Kdm6a* cKO testes (**Fig 2D and 2E**). Gamma-H2AX, a phosphorylated histone variant induced at sites of DNA damage and involved in meiotic recombination, meiotic sex chromosome inactivation and meiotic silencing of unsynapsed chromatin, also exhibited similar cellular distribution patterns and global levels in *Kdm6a* cKO testes relative to control (**Fig 2E and 2F**), implying that these processes progress normally in the absence of KDM6A.

**Fig 2.**
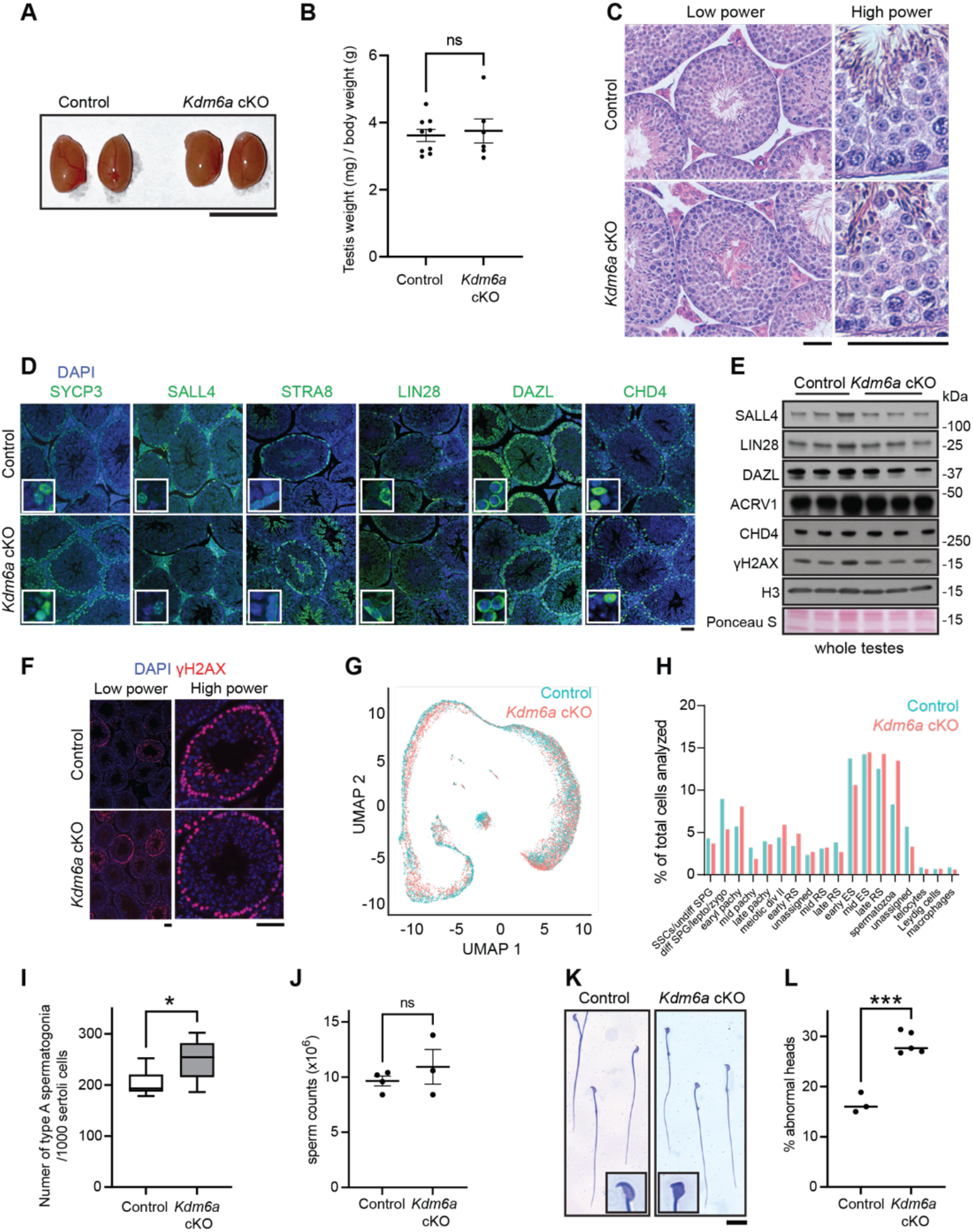
Minimal impact on spermatogenesis in *Kdm6a* cKO mice. (**A**) Gross morphology of testes from a control and a *Kdm6a* cKO mouse. Scale bar = 1 cm. (**B**) Testis weights from control and *Kdm6a* cKO mice normalized to body weight. (**C**) Cross-sections of control and *Kdm6a* cKO seminiferous tubules stained with hematoxylin and eosin. Scale bars = 50 μm. (**D**) Control and *Kdm6a* cKO testis sections immunostained (green) for spermatogonia markers (LIN28A, STRA8, CHD4, and SALL4), a spermatocyte marker (SYCP3), and a broad germ cell marker (DAZL) as well as DAPI to stain DNA (blue). Scale bar = 50μm. (**E**) Whole lysates from control and *Kdm6a* cKO testes (n = 3) immunoblotted for spermatogonia markers (LIN28A, SALL4 and CHD4), a round spermatid marker (ACRV1), and a marker for meiotic progression (γH2AX). Histone H3 and ponceau S. serve as loading controls. (**F**) Immunofluorescence staining (red) for γH2AX in seminiferous tubules of control and *Kdm6a* cKO mice. (**G**) UMAP plot of single cell RNA-seq data from control (blue) and *Kdm6a* cKO testis (pink). (**H**) The percentage of cells classified into each cluster out of total cells profiled in control and *Kdm6a* cKO scRNA-seq datasets. (**I**) Quantification of type A spermatogonia by visual examination of testes sections stained with hematoxylin and eosin. (**J**) Epididymal sperm counts after swim-out in control and *Kdm6a* cKO mice. (**K**) Coomassie-stained spermatozoa from control and *Kdm6a* cKO cauda epididymides. Scale bar = 20μm. (**L**) Manual blinded quantification of abnormal heads detected by brightfield microscopy of Coomassie-stained sperm from control and *Kdm6a* cKO mice.

We next generated an scRNA-seq dataset from *Kdm6a* cKO testes and asked if *Kdm6a* loss leads to changes in the relative numbers of specific spermatogenic cell types compared to our control data set (see **Fig 1C**). We found similar proportions of all cell types between control and *Kdm6a* cKO testes (**Fig 2G and 2H**). No significant difference in the number of preleptotene spermatocytes was detected by manual counting in hematoxylin and eosin stained testis sections (**Fig S2C**). We did detect a statistically significant but very modest increase in the numbers of type A spermatogonia in *Kdm6a* cKO testis sections relative to control (**Fig 2I**).

Finally, we explored the possibility that loss of *Kdm6a* affects spermiogenesis and alters the form or function of mature spermatozoa. The number of sperm in the cauda epididymides were unaffected (**Fig 2J**). There was a modest but statistically significant increase in the fraction of *Kdm6a* cKO sperm that exhibited triangular shaped head defects (29%) relative to control (17%, **Fig 2K and 2I**). No motility defect was detected in *Kdm6a* cKO sperm under uncapacitated or capacitated conditions using computer-assisted semen analysis (CASA, **Fig S2D**).

We conclude that there is no meaningful spermatogenesis defect in *Kdm6a* cKO mice and that KDM6A is dispensable for sperm production and fertility.

### KDM6A does not significantly impact chromatin accessibility in spermatogonia

KDM6A has been shown to regulate chromatin accessibility in several mammalian cell types, primarily through H3K27me3-independent mechanisms (33-35). We asked if KDM6A similarly contributes to chromatin remodeling in the male germline. Western blotting in subcellular fractions of whole testis confirmed that as expected, KDM6A protein is predominantly nuclear, indicating that the bulk of its activity is likely directed towards chromatin targets in the male germ line (**Fig S3A**). We therefore interrogated genome-wide changes in chromatin accessibility provoked by KDM6A loss in the male germline by performing ATAC-seq in sorted cKIT+ cells from *Kdm6a* cKO and control mice. The average number of peaks called was similar between control (n = 26,184) and *Kdm6a* cKO samples (n = 24,671). Peak overlap between replicates was high for both control and *Kdm6a* cKO samples, although there was more difference between cKO samples likely owing to variabilities inherent in cell sorting procedures (**Fig 3A, S1 Dataset)**. Strong ATAC-seq peaks were predominantly identified at gene promoters as expected (**Fig S3B, S3C, and S3D**). Differentially accessible regions (DAs) between control and *Kdm6a* cKO samples were called at *P* < 0.01 using the *csaw* pipeline with loess normalization (36). We detected only 24 DAs that gained chromatin accessibility and 12 DAs that lost chromatin accessibility in *Kdm6a* cKO cKIT+ cells relative to control (**Fig 3B**). These changes were modest in magnitude and frequently occurred outside of promoters for gained DAs (**Fig 3C**).

**Fig 3.**
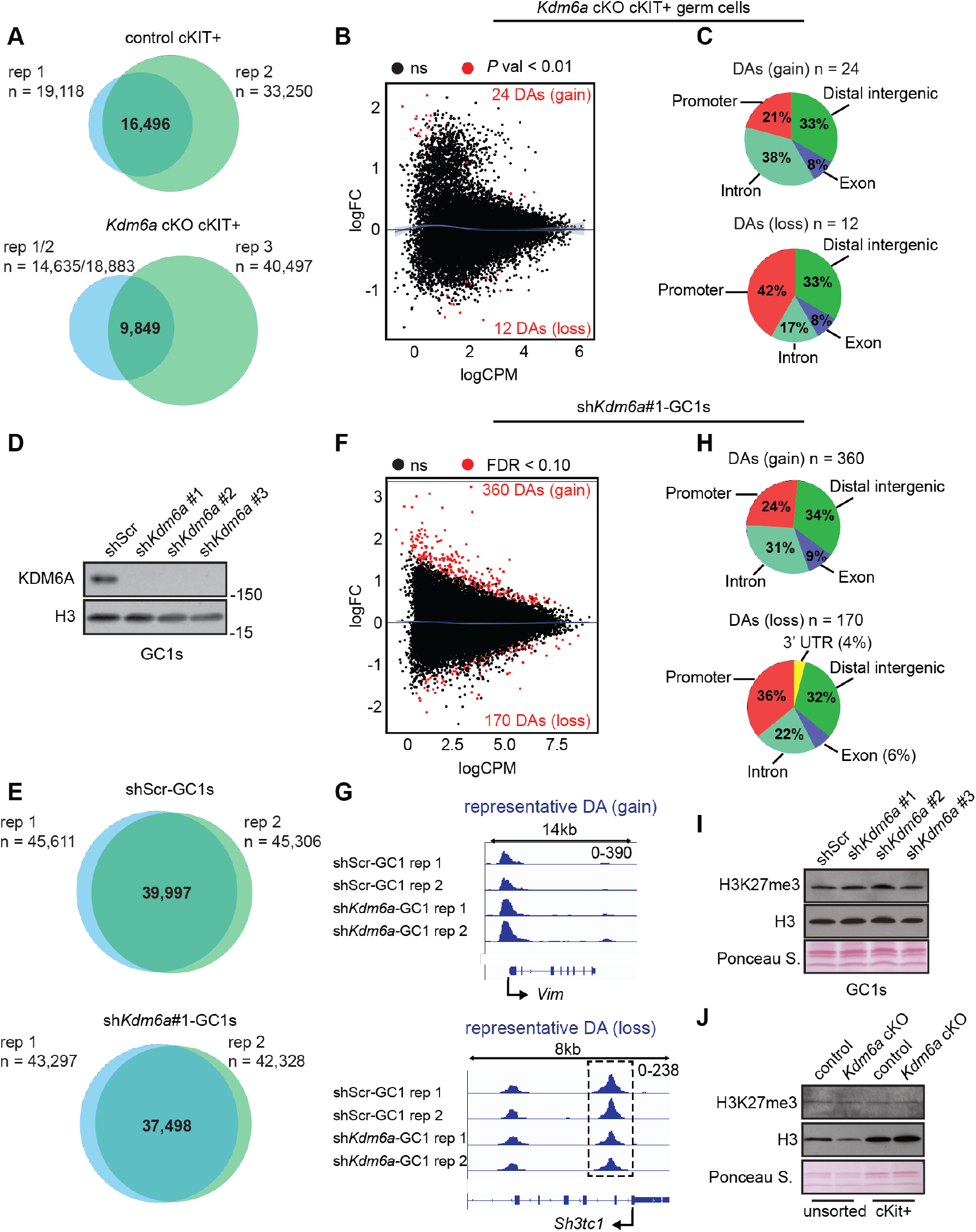
Male germ cells depleted for KDM6A show modest changes in chromatin accessibility. (**A**) Intersection of genomic coordinates for ATAC-seq peaks across replicates of cKIT-sorted cells from control and *Kdm6a* cKO testes. (**B**) MA plot for ATAC-enriched regions showing the distribution of differential chromatin accessibility in *Kdm6a* cKO cKIT+ cells. Black dots represent non-significant regions (ns) and red dots represent significant differentially accessible (DA) regions (p <0.01). Blue lines are loess fits to each distribution. (**C**) Gene feature distributions for DA regions detected in *Kdm6a* cKO cKIT+ cells. (**D**) Western blotting for KDM6A and histone H3 (loading control) in GC1-SPG cells expressing shScramble (shScr) or one of three different short-hairpin RNAs targeting *Kdm6a* mRNA. (**E**) Intersection of genomic coordinates for ATAC-seq peaks across replicates of shScr-GC1-SPGs and sh*Kdm6a*-GC1-SPGs. (**F**) MA plot for ATAC-enriched regions showing the distribution of differential chromatin accessibility in shKdm6a-GC1-SPGs. (**G**) Genome browser tracks for representative regions of chromatin accessibility gain (above) and loss (below) in sh*Kdm6a*-GC1-SPGs. (**H**) Gene feature distributions for DAs detected in sh*Kdm6a*-GC1-SPGs. (**I**) Western blot for H3K27me3 and histone H3 in shScr-GC1-SPGs and sh*Kdm6a*-GC1-SPGs. (**J**) Western blot for H3K27me3 and histone H3 in unsorted and cKIT-sorted cells from control and *Kdm6a* cKO testes.

Because variability in cell sorting limited our ability to detect subtle changes in accessibility in spermatogonia in vivo, we turned to a more controlled in vitro system, GC1 spermatogonial-derived cells, in order to minimize sample variability and enable robust quantitative comparisons. Expressing either of two short hairpin RNAs to *Kdm6a* in GC1 cells robustly depleted *Kdm6a* at the protein level (**Fig 3D**). ATAC-seq in GC1 cells expressing short hairpins against *Kdm6a* (sh*Kdm6a*) or scrambled control (shScr) yielded an average of 44,135 peaks across all samples. Overlap of peak coordinates across replicates was high for both sh*Kdm6a* and shScr (**Fig 3E, S1 Dataset**), and peaks in both groups were predominantly localized to promoters of transcriptionally active genes (**Fig S3E, S3F and S3G**). Notably, we also found that approximately 70% of peak coordinates defined in cKIT-sorted cells from control testes intersected those called in shScr-GC1 cells, indicating that accessibility in GC1 cells is broadly similar to spermatogonia in vivo (**Fig S3H**). In contrast to the paucity of DAs detected in cKIT+ cells, we identified 530 highly significant DAs even with a more stringent threshold (FDR < 0.10) in sh*Kdm6a*-GC1 cells (**Fig 3F and 3G**). Most DAs represented a gain in chromatin accessibility contrasting with the expected role of KDM6A in promoting accessibility. However, the DA regions that lost accessibility were more likely to occur at promoter regions, potentially indicating a more canonical and/or more direct activity of KDM6A at promoters (**Fig 3H**). While the pattern of gene feature distributions for gained and lost DAs was similar between cKIT-sorted *Kdm6a* cKO cells and sh*Kdm6a*-GC1 cells, there were no DAs called in common between the two cell types.

Because KDM6A is a H3K27me3 demethylase, we asked if cKIT+ sorted *Kdm6a* cKO cells or sh*Kdm6a-*GC1 cells exhibited global gains in H3K27me3 relative to control cells to support a link between detected DAs and histone modification. We found that the global levels of H3K27me3 as assayed by Western blotting were comparable between control lysates and KDM6A-depleted lysates (**Fig 3I and 3J**). These findings imply that any changes to H3K27me3 are locus-specific, as previously seen in sperm of *Kdm6a* cKO mice (8), or that KDM6A mediates chromatin remodeling via regulation of alternative modifications, such as H3K27ac or H3K4me1.

Finally, we compared our accessibility data in cKIT+ sorted spermatogenic cells with differentially methylated regions (DMRs) previously identified in sperm of *Kdm6a* cKO animals (8). Five hundred thirty-nine (6%) of all DMRs overlap regions of open chromatin in spermatogonia, a six-fold enrichment over the amount of overlap observed when DMR intervals were randomly shuffled across the genome (*P* < 0.0001, z-test for one proportion). Thus, chromatin accessibility in *Kdm6a*-expressing spermatogonia is modestly but statistically significantly associated with sites of altered DNA methylation in sperm following KDM6A depletion.

Together, these results show that KDM6A plays a limited role in regulating chromatin accessibility in differentiating spermatogonia in vivo.

### KDM6A is a transcriptional activator in pre-meiotic male germ cells

KDM6A is a positive regulator of transcription in mammalian cells through its H3K27 demethylase activity and its association with MLL3/4 complexes to promote H3K4 mono-methylation at promoters and enhancers (10-12, 37). We next sought to define any genome-wide transcriptional effects of *Kdm6a* loss in the male germline by calling differentially expressed genes (DEGs) between *Kdm6a* cKO and control testes from our scRNA-seq datasets. Across all clusters, a total of 886 distinct genes exhibit significantly altered expression (**S2 Dataset**), with the greatest number of DEGs found in cells where *Kdm6a* is most highly expressed and detectable as a DEG (**Fig 4A, S4A**). To validate these data, we also performed bulk RNA-seq in control and *Kdm6a* cKO whole testes (n=3) and confirmed using gene set enrichment analysis (GSEA, (38, 39)) that similar sets of genes exhibited altered expression (**Fig S4B**). Deregulation of gene expression was strongly biased towards downregulation (76% of DEGs, **Fig 4B**), consistent with the known function of *Kdm6a* as a transcriptional activator. Upregulated DEGs are enriched for Gene Ontology (GO) terms related to cytoplasmic translation, while downregulated DEGs were more weakly enriched for several germ-cell related terms, the most significant being “spermatid development” (**Fig 4C**). Notably, while the expression of the homologs *Uty* and *Kdm6b* was unchanged, we found that other histone lysine demethylases (*Kdm1a, Kdm3a, Kdm5a, Kdm5b, Kdm3c*) and chromatin remodelers (*Smarca2* and *Chd5*) were downregulated in *Kdm6a* cKO testes (**Fig 4D**). Based on the strong bias toward downregulation and known role of KDM6A as a transcriptional activator, we focused our subsequent analysis on the downregulated DEGs detected in *Kdm6a*-expressing cell populations (i.e., clusters 4 and 5) as they are most likely to represent direct target genes of KDM6A. These high-confidence DEGs were enriched for GO terms related to meiosis, sperm motility, histone modification and chromosome organization (**Fig 4E**). Interestingly, we found that approximately half of DEGs identified in cluster 3, representing the terminal stage of spermatogenesis (fully elongated and condensed sperm heads), were also called as DEGs in either cluster 4 or cluster 5 (**Fig 4F**). A smaller subset of these genes (n = 25), almost all of which are downregulated, was identified as DEGs across all three clusters (**Fig 4G**), suggesting that for some broadly expressed genes, KDM6A establishes regulatory information that persists across spermatogenesis.

**Fig 4.**
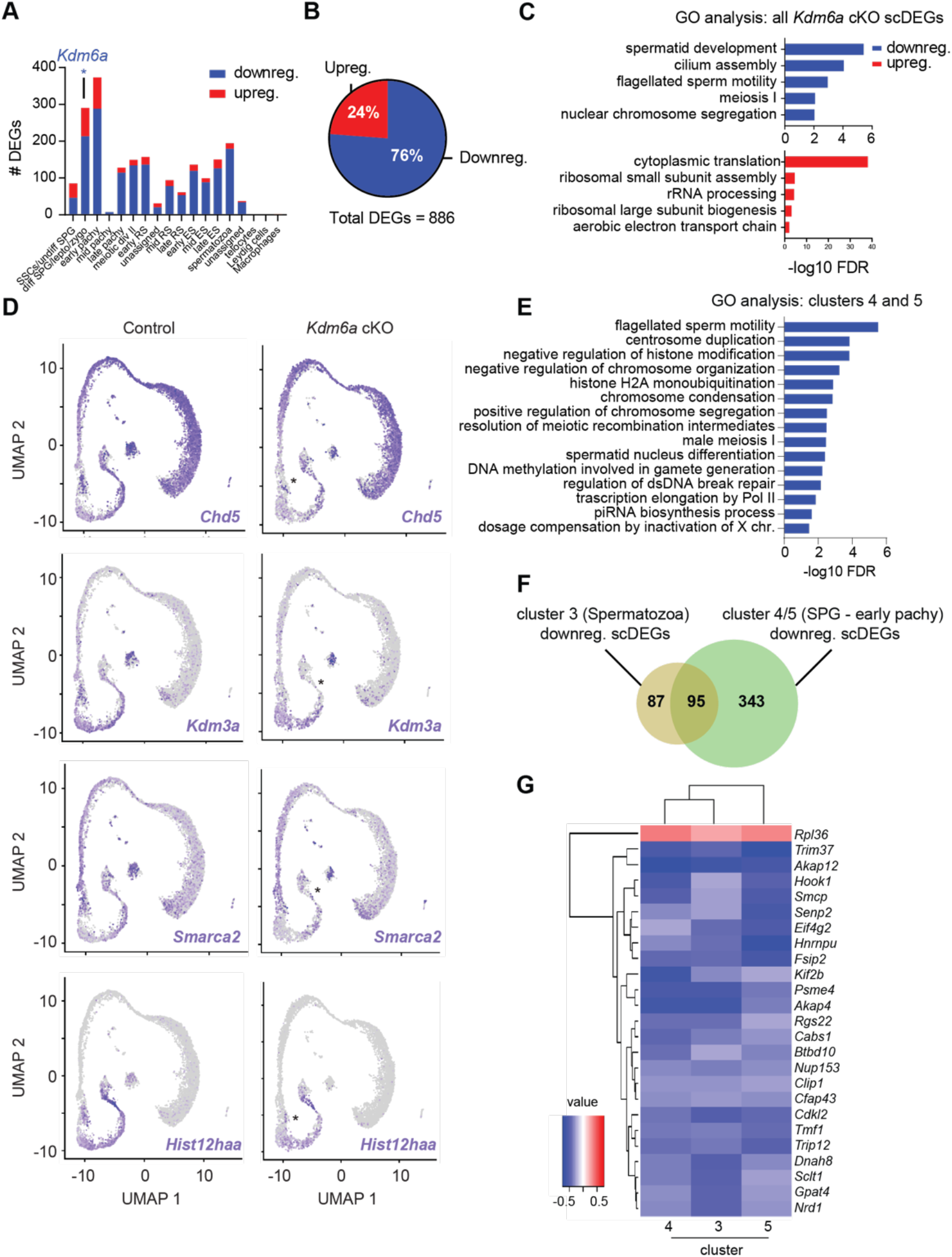
KDM6A is a Transcriptional Activator in Early Spermatogenic Cells. (**A**) Fraction of differentially expressed genes identified by scRNA-seq (scDEGs) that are downregulated and upregulated in *Kdm6a* cKO testis relative to control. (**B**) Numbers of scDEGs identified per cluster, labeled by testis cell identity. *Kdm6a* is identified as a scDEG in cluster 4, which comprises differentiated spermatogonia and early stages of meiotic prophase. (**C**) Gene ontology analysis showing terms enriched in all downregulated and upregulated scDEGs identified in *Kdm6a* cKO testes. (**D**) UMAPs showing the expression for representative downregulated scDEGs associated with chromatin organization and remodeling. Asterisks (*) mark cells with most profound changes in expression. (**E**) Gene ontology terms enriched in scDEGs identified for clusters 4 and cluster 5 i.e., cell populations spanning differentiated spermatogonia to early pachytene. (**F**) Overlap of scDEGs identified in cells classified into cluster 3 (spermatozoa) and clusters 4/5 (differentiating spermatogonia and early meiotic cells). (**G**) Heatmap showing the normalized gene expression levels in *Kdm6a* cKO testis for 25 genes that are deregulated in cluster 3, cluster 4, and cluster 5.

We did not find an association between genes near DAs called in cKIT+ sorted cells from *Kdm6a* cKO testes (see previous section) and DEGs identified in our scRNA-seq dataset, likely due to the variability in the ATAC-seq data derived from sorted cKIT+ cells. We therefore applied our GC1 knockdown model to ask if transcriptional and chromatin accessibility effects of KDM6A depletion were correlated in GC1 cells. We assessed expression changes between sh*Kdm6a* and shScr GC1 cells and detected 1,463 DEGs (*P* adj < 0.05, **S3 Dataset**) in sh*Kdm6a#1*-GC1 cells relative to shScr control cells, with *Kdm6a* being one of the most significantly downregulated genes (**Fig S4C**). Consistent with our in vivo findings, there was a bias towards downregulation (77% of total DEGs *P* adj < 0.05, > 1.5 fold change). Expression of a different short hairpin RNA sequence to *Kdm6a* (*shKdm6a#*2) yielded similar results, confirming that the observed gene expression changes were not due to off-target effects (**Fig S4D, S4E, S3 Dataset**). GSEA revealed a significant enrichment for genes associated with loss of chromatin accessibility in sh*Kdm6a*#1-GC1 cells and downregulated genes (**Fig S4F**). We did not find any enrichment for genes associated with gains in chromatin accessibility and upregulated gene expression. Only twenty-four DEGs detected in sh*Kdm6a*#1-GC1 cells were also called in our *Kdm6a* cKO scRNA-seq dataset (**Fig S4G**). Interestingly however, these included several genes important for spermatogenesis, such as *Chd5*, a master regulator of the histone-to-protamine transition (40) and *Cdc14a*, an essential factor for male fertility in mouse and human (41).

We conclude that KDM6A promotes timely expression of genes related to chromatin organization predominantly during meiotic entry in the male germline. Loss of KDM6A leads to altered transcriptional states during meiotic entry as well as later in spermatogenesis. There is little association between transcriptional effects and KDM6A-directed chromatin remodeling, although some subtle transcriptional changes detected in our in vitro model may be mediated by KDM6A-directed remodeling events.

### A subset of *Kdm6a* cKO DEGs are persistently deregulated in the germlines of wildtype offspring

The soma of wild type mice derived from a paternal germline lacking KDM6A (*Kdm6a* F1) exhibit transcriptomic and DNA methylation changes (8). These changes were reported to be greater in magnitude in the soma of wildtype mice that were derived from two successive generations of paternal germline lacking KDM6A (*Kdm6a* F2, **Fig 5A**), implying that KDM6A activity in the male germline confers gene regulatory information important for the next generation that is additively perturbed in its absence over successive generations. We therefore asked if gene expression changes could be detected in the male germ lines of *Kdm6a* F1 and *Kdm6a* F2 mice despite expression of functional *Kdm6a* (**Fig S5A**).

**Fig 5.**
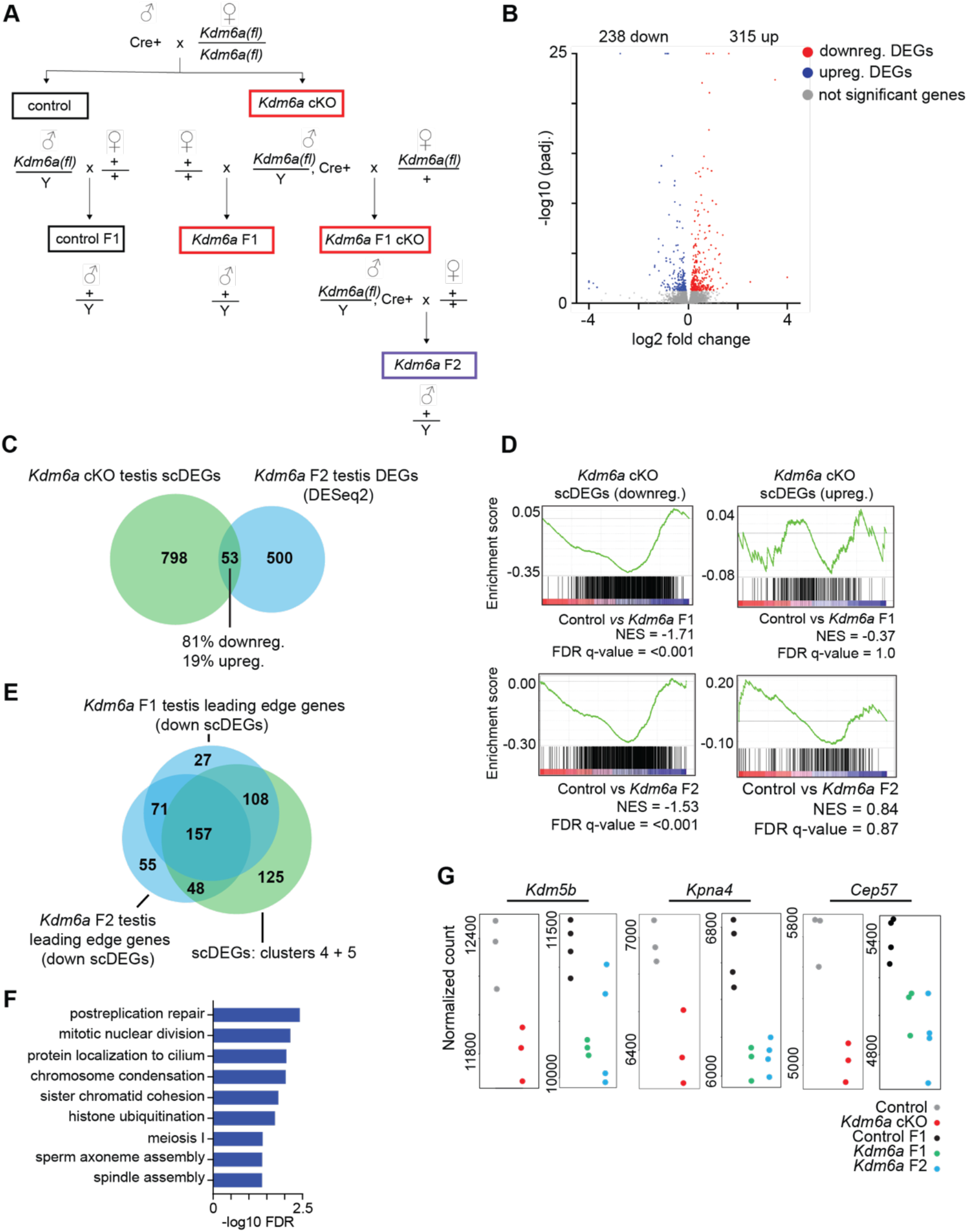
A subset of differentially expressed genes in the *Kdm6a* cKO germline is persistently deregulated in testes of wildtype offspring. (**A**) Breeding scheme for the generation of *Kdm6a* cKO mice and wildtype progeny: *Kdm6a* F1 and *Kdm6a* F2. (**B**) Volcano plot showing the gene expression changes in *Kdm6a* F2 testes relative to control. (**C**) Overlap of differentially expressed genes (DEGs) detected by single cell RNA-seq (scRNA-seq) in *Kdm6a* cKO testis and DEGs identified by analysis of bulk RNA-seq of *Kdm6a* F2 teste**s**. (**D**) Gene set enrichment analysis (GSEA) plots showing the enrichment of *Kdm6a* F1 (top) and *Kdm6a* F2 testis (bottom) expression data for all single cell DEGs (scDEGs) identified in *Kdm6a* cKO testis. The non-significant enrichment for upregulated scDEGs is shown by the GSEA plots to the right. Normalized enrichment score = NES. (**E**) Overlap of scDEGs for clusters 4 and 5 in *Kdm6a* cKO testis with the leading-edge genes (LEGs) identified by the GSEA of *Kdm6a* F1 and *Kdm6a* F2 testis expression data. (**F**) Gene ontology terms enriched for leading edge genes shared between *Kdm6a* F1 and *Kdm6a* F2 gene expression datasets. (**G**) Normalized counts plotted for representative DEGs that are persistently deregulated across the germline of *Kdm6a* cKO, *Kdm6a* F1, and *Kdm6a* F2 mice.

Bulk RNA-seq analysis of whole adult testis data from *Kdm6a* F1 mice identified only sixty-four DEGs (**S4 Dataset**), indicating that the overall germ cell transcriptome is largely unaffected by KDM6A loss in the paternal germline. However, we identified 531 DEGs (*P* adj. <0.05, **S4 Dataset**) in whole adult testes of *Kdm6a* F2 mice, implying that absence of KDM6A in the male germ line for two successive generations results in cumulative changes in the gene regulatory state of the male germline in offspring (**Fig 5B**). Unlike the *Kdm6a* cKO germline, we did not detect any bias in the direction of deregulation. Fifty-three DEGs (10% of all *Kdm6a* F2 DEGs) detected in *Kdm6a* F2 testes were also identified as DEGs in our scRNA-seq dataset from *Kdm6a* cKO testis with the bulk of this overlap coming from downregulated DEGs (**Fig 5C**). This result suggested that a subset of DEGs induced by loss of *Kdm6a* in the parental germline is persistently deregulated across generations. GSEA with *Kdm6a* F1 and F2 whole testis expression data revealed a significant (nominal *P* < 0.0001) enrichment for downregulated, but not upregulated, scDEGs identified in *Kdm6a* cKO testis (**Fig 5D**). We found a high overlap between the genes that contributed most to this enrichment signal (‘leading-edge genes’) in *Kdm6a* F1 and *Kdm6a* F2 datasets, indicating that a defined subset of genes is persistently, albeit mildly, deregulated across generations (**Fig 5E**). GO analysis revealed that this subset of genes is predominantly enriched for terms related to chromosome organization and histone modification (**Fig 5F**); notable examples included the chromatin modifiers *Chd5* and *Kdm5b*. The majority of leading-edge genes (∼69%) were identified as downregulated scDEGs in cluster 4 and cluster 5, implying that most persistently deregulated genes represent those of putative direct KDM6A target genes in the male germline (**Fig 5E**). Consistent with this finding, GSEA of *Kdm6a* F1 and *Kdm6a* F2 expression datasets revealed a greater enrichment for downregulated scDEGs identified in cluster 4 and cluster 5 relative to scDEGs for all other clusters (**Fig S5B and S5C**). We conclude that KDM6A activity in the paternal germline contributes to gene regulatory state in the germlines of male offspring.

## Discussion

Mice derived from a paternal germline lacking *Kdm6a* have a reduced lifespan and more readily develop cancers (8). Thus, KDM6A likely has activities in the male germline that are important for gene regulation in the next generation. To identify these activities and begin to understand how they relate to offspring health, we evaluated the regulatory and phenotypic consequences of deleting *Kdm6a* in the postnatal male germ line. We found that *Kdm6a* is expressed specifically and transiently in late spermatogonia and during early meiosis, and that while KDM6A loss is compatible with normal sperm production and fertility, it perturbs gene regulatory programs that have lasting effects in late spermatogenesis and in germ cells of subsequent generations.

Although previous studies reported broad KDM6A expression across adult human and mouse seminiferous epithelium, including spermatocytes and round spermatids (42, 43), we found that *Kdm6a* expression was limited to late spermatogonia and the early stages of meiotic prophase. This discrepancy could be explained by cross-reactivity of anti-KDM6A antibodies in tissue sections; indeed, we excluded several commercially-sourced antibodies from our study due to non-specific binding in cKO testis sections. Given its precisely timed expression, we speculate that KDM6A may participate in balancing the deposition of histone modifications during the mitosis-meiosis transition, and that rapid downregulation of KDM6A expression soon after early prophase is important for maintaining an appropriate histone methylome at this key developmental timepoint. Histone methylation plays a critical role during meiotic entry: histone methyltransferases participate in meiotic recombination, synapsis, and chromosome segregation (44), and major changes in histone methylation, including accumulation of H3K27me3, occur on sex chromosomes during prophase I to mediate transcriptional silencing (45). Our scRNA-seq data highlights KDM6A as a transcriptional activator that supports the meiotic gene expression program, albeit in a manner dispensable for meiotic progression overall. Along with mESCs, germ cells are also uniquely rich in bivalent promoters co-occupied by H3K27me3 and H3K4me3. The bivalent state facilitates the mitosis-meiosis transition (46) and is thought to encode regulatory information for embryonic development (25). KDM6A has been shown to resolve bivalent domains to univalent states during differentiation of mESCs and human neural progenitor cells (18, 47). Future experiments in which KDM6A is ectopically expressed in post-meiotic germ cells will be of interest to better determine the significance of its restricted developmental expression.

While loss of KDM6A has virtually no effect on spermatogenic phenotype, we found that it has a substantial effect on gene expression. The majority of changes in gene expression occurred in the same cell populations that normally express KDM6A, but many expression changes also occurred during late spermatogenesis, implying that regulatory consequences of KDM6A loss extend throughout spermatogenic development. DEGs identified in the *Kdm6a* cKO were enriched for genes encoding other epigenetic modifiers, suggesting that the gene regulatory activity of KDM6A can indirectly affect the chromatin landscape in the male germline. For example, regulators of histone ubiquitination represent a major subset of DEGs following KDM6A loss, making changes in the global distribution of this modification in the *Kdm6a*-null germline a promising question for future studies. Additionally, KDM6A may be active towards other histone substrates besides H3K27me3, including the enhancer marks H3K4me1 and H3K27ac (33, 37). We previously observed locus-specific changes in levels of H3K27me3 in mature spermatozoa from *Kdm6a*-null mice corresponding to redistribution of the H3K27me3 modification (8). Further work defining the genome-wide distribution of H3K27me3, H2AK119ub, H3K4me1, and H3K27ac throughout spermatogenesis in the *Kdm6a* cKO will help to address the extent to which KDM6A indirectly affects gene expression via modulation of the expression levels of other histone modifiers. Low-input methods can also be used to determine genome-wide occupancy of KDM6A in early spermatogenic cells in order to identify its direct targets.

KDM6A loss has been shown to have strong effects on chromatin accessibility and the expression of corresponding genes in other cell types (33, 34), but we did not find a similarly strong effect on accessibility in the male germline. This could be because the population of late spermatogonia we were able to sort based on cKIT marker expression does not precisely correspond to the window of KDM6A expression during spermatogenesis. In GC1 cells depleted for KDM6A, we identified more, stronger changes in accessibility compared to cKIT-sorted cells, but we did not find a bias toward loss of accessibility as expected based on the known role of KDM6A as a transcriptional activator, nor did we find a strong connection to gene expression changes. This result is consistent with a previous ATAC-seq study in *Kdm6a*-null hematopoietic stem cells which revealed an approximately equal balance between gain and loss of chromatin accessibility and little correlation with KDM6A occupancy at corresponding genes. Further work is needed to understand if KDM6A can directly promote chromatin accessibility or if its accessibility effects are primarily indirect.

Based on the lack of a spermatogenic phenotype in *Kdm6a* cKO mice, we considered the possibility of redundancy with KDM6B and UTY, homologs that have been shown to function redundantly during mouse embryonic development and cellular differentiation, respectively (48, 49). *Uty* transcript abundance is very low in testes, only partly overlaps in developmental timing with *Kdm6a*, and is not upregulated in the context of the *Kdm6a* cKO, leading us to conclude that *Uty* likely does not compensate for *Kdm6a*. KDM6B expression does overlap with KDM6A and the phenotype of *Kdm6b* germline cKO males is similar: *Kdm6b* cKO males are fertile with higher numbers of undifferentiated spermatogonia (42). Therefore, *Kdm6b* may be partially redundant with *Kdm6a* during spermatogenesis. The lack of global changes in H3K27me3 we observed in *Kdm6a* cKO testes mirrors previous reports in the *Kdm6b*-null male mouse germline and further supports redundancy between these homologs in spermatogenesis. KDM7B may also have some functional redundancy with KDM6A given that their expression profiles are very similar across spermatogenesis. Like *Kdm6a, Kdm7b* is X-linked in mammals and can demethylate H3K27 in zebrafish (31). Definitive assessment of redundancies between KDM6A and UTY, KDM6B, and KDM7B in the male germline will require generation of mice multiply knocked out for each of these factors.

All together, our analysis of spermatogenesis in *Kdm6a* cKO mice excludes sperm dysfunction as a primary explanation for the previously observed intergenerational effect on lifespan and cancer. Instead, we provide evidence supporting an intergenerational role for KDM6A activity in regulation of gene expression in the germ line by showing that some expression changes induced by KDM6A loss in spermatogenic cells are retained in the germ cells of subsequent generations. This observation is surprising, since acquired epigenetic information is largely reset in the next generation both soon after fertilization and during primordial germ cell development. Our findings here mirror a recent report in *C. elegans* showing that the status of H3K27me3 at sperm alleles is inherited across generations to regulate gene expression in the germline (50). A similar mechanism may be at play in our mammalian model given that KDM6A targets H3K27me3. We speculate that loss of KDM6A in the male germline permits accumulation of epimutations, some of which are resistant to erasure in the next generation and are difficult to reset even in the presence of a functional *Kdm6a* allele. Therefore, KDM6A may act during the dynamic chromatin reorganization that occurs at the start of meiosis to safeguard the male germline from acquiring altered epigenetic states (**Fig 6**). Future work will define the nature and location of these epimutations, as well as the mechanism by which they might resist reprogramming.

**Fig 6.**
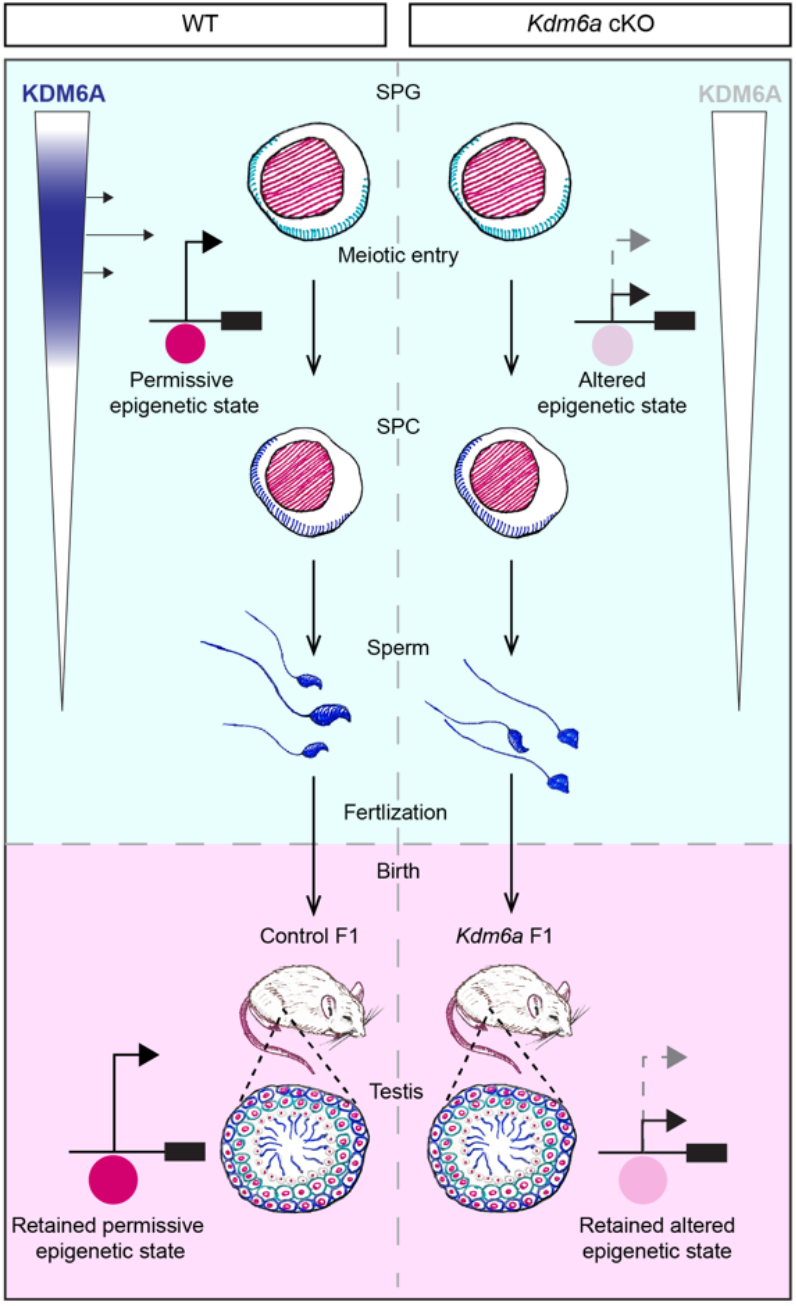
Model for intergenerational inheritance of gene regulatory states following deletion of *Kdm6a* in the male germline. KDM6A is predominantly expressed during the mitotic-to-meiotic transition in the male germline where it functions as a transcriptional activator putatively through promoting permissive epigenetic states. Absence of KDM6A may alter epigenetic states to perturb gene expression. These altered gene regulatory states in *Kdm6a* cKO sperm resist reprogramming at a subset of loci upon feralization and development, resulting in perturbed transcriptional profiles in the *Kdm6a* F1 germline. Spermatogonia (SPG) and Spermatocyte (SPC).

## Materials and Methods

### Mouse breeding and animal care

All mice were maintained on a C57BL/6J genetic background. To obtain *Kdm6a* cKO males, *Kdm6a*^flox/flox^ (*Kdm6a*^*tmc1(EUCOMM)Jae*^) females were mated with mice carrying one copy of the *Ddx4*-Cre transgene (B6-*Ddx4*^*tm1*.*1(cre/mOrange)Dcp*^) (51, 52) in which Cre is expressed specifically in germ cells beginning at embryonic day 15.5. *Kdm6a* F1 males were generated by crossing male *Kdm6a* cKO mice with wildtype females. *Kdm6a* F1 cKO males were generated by breeding male *Kdm6a* cKO mice with female *Kdm6a*^*flox/+*^ mice, which were then crossed with wildtype females to generate wildtype *Kdm6a* F2 males. Primers used for genotyping are listed in **Table S1**. These studies were approved by the Yale University institutional animal case and use committee under protocol 2020-20169. All mice used in these studies were maintained and euthanized according to the principles and procedures described in the National Institutes of Health guide for the care and use of laboratory animals.

### GC1-SPG culture and transduction

GC-1spg cells (ATCC, CRL-2053) were cultured in DMEM (Gibco, 11965092) supplemented with 10% fetal bovine serum (Corning, 35-011-CV), 1 mM sodium pyruvate (Gibco, 11360-070), and pen-strep (Gibco, 15140-122). At 90% confluency, cells were passaged 1:15 every 3 days using trypsin (Gibco, 25300054) and centrifugations at 200 xg. *Kdm6a* knockdown cell lines were generated by reverse lentiviral transduction of shRNAs expressed from a pZIP-mEF1α-ZsGreen-Puro lentiviral vector (TransOMIC Technologies) in the presence of 8 μg/ml polybrene. Three days after transduction, the cells were treated with 2 μg/ml puromycin for a week to select for stable transductants. Sequences for shRNA constructs can be found in **Supplementary Table S1**. MycoStrip™ (InvivoGen, rep-mys-10) did not detect mycoplasma contamination in any of the cell cultures used in this study.

### Immunoblotting

Whole cell lysates were prepared using RIPA buffer supplemented with 0.9% SDS. Subcellar fractions were prepared according to Gagnon *et al*. (2014). Up to 40μg of total protein was loaded onto a Mini-PROTEAN TGX gel (Bio-Rad, 456-8093) and separated by SDS-PAGE for 1 hour at 200 volts. Separated proteins were then wet transferred to a 0.45 μm nitrocellulose membrane (GE Healthcare Life Science, 10600003) in Towbin buffer (25mM Trizma base, 192 glycine, 20% (v/v) methanol) at a constant current of 250mA for 1 hour. The membrane was then briefly incubated in Ponceau S (Sigma Aldrich, P7170), washed in TBST, then blocked in 5% w/v non-fat dry milk (American Bio, ab10109-01000) in TBST for 30 mins with gentle agitation at room temperature. Primary antibodies were diluted in blocking buffer and incubated with the membrane overnight at 4°C with gentle rocking. After washing in TBST the membranes were incubated with peroxidase conjugated secondary antibodies diluted 1:10,000 in blocking buffer. Following washing, the membrane was incubated for 5 mins in SuperSignal™ West Pico Plus (Thermo, 34580) or the Femto variant (Thermo, 34096) and the signal was captured with X-ray film (Thermo, 34090). Antibodies used for immunoblotting are listed in **Table S2**.

### Tissue staining

Mouse testes were dissected and immersed in Hartman’s fixative for 30 mins at room temperature before bisecting for further fixation overnight at 4°C on an end-to-end rotator. Fixed testes were subjected to standard dehydration and clearing processing for paraffin embedding. Wax sections were stained with hematoxylin and eosin for histological analysis and indirect immunofluorescence was performed after heat-induced epitope retrieval with citrate buffer (Vector Laboratories, H-3300). Confocal images were taken using a Zeiss LSM 880 microscope. Antibodies used for immunostaining are listed in **Table S2**. In situ hybridization was performed using RNAscope 2.5 HD assay-RED (ACDBio, 322360) according to the manufacturer’s instructions. The assay was optimized by performing heat-induced antigen retrieval for 15 mins and protease III digestion for 30 mins at room temperature.

### Immunocytochemistry

Meiotic spreads were prepared as previously described (54) for subsequent indirect immunofluorescence. Confocal images were taken using a Zeiss LSM 880 microscope. Antibodies used for staining are listed in **Table S2**.

### Testis cell dissociation

Decapsulated testes were incubated in DMEM containing 0.75 mg/ml collagenase type IV (Gibco, 17104-019) for 10 mins at 37°C with occasional inversion. Equal volume of DMEM was added to the suspension of dissociated tubules before centrifuging at 400 xg for 5 mins at 4 °C. After removing the supernatant, the cell pellet was washed in DMEM and resuspended in 0.05% trypsin-EDTA (Gibco, 25200-056) for 10 mins incubation at room temperature. An equal volume of ice-cold DMEM+10%FBS was then added to the cell suspension before washing the pellet twice with media. The cell pellet was resuspended in DPBS (Sigma, D8537) containing 0.04% bovine serum albumin (sigma, A9647) and filtered through a 100 μm filter (Falcon, 22363549) then a 40 μm filter (Falcon, 352340) to generate a single cell suspension.

### Spermatogonia enrichment by flow cytometry

Dissociated germ cells were incubated in the dark for 30 mins with PE conjugated anti-cKIT at 1:800 (Thermo, 12-1171-82). Stained cells were then washed twice, resuspended to 10 million cells/ml and sorted on a two-laser (466 nm and 561 nm) cell sorter (Bio-Rad S3) as previously described (55). Briefly, debris was removed from the plot based on FSC-A/SSC-A and singlets were selected based on FSC-H/FSC-A. A histogram of FL1-A was used to identify positively stained cells compared to an unstained aliquot.

### Morphological assessment of spermatozoa

Cauda epididymides were dissected from adult male mice and cut into 1 ml of warm M2 medium (Sigma, M7167) to allow sperm to swim out. After 15 mins, the cell suspension was centrifuged for 2 mins at 2000 xg at 4 °C and the resulting pellet was resuspended in 4% paraformaldehyde for 20 mins at room temperature. Fixed spermatozoa were pelleted and washed twice in 100 mM ammonium acetate before resuspending in 150 μl wash buffer. 5 μl of this cell suspension was spread across a glass slide and air dried followed by Coomassie staining for 10 mins. After rinsing in tap water, the slides were air dried then coverslipped with DPX (Sigma, 06522). The percentage of sperm with abnormal heads was visually assessed from >400 sperm for each biological replicate using brightfield microscopy under blinded conditions.

### Motility assessment of spermatozoa

Cauda epididymal sperm were collected by swim-out (described above) and aliquots were placed in a slide chamber (Cell Vision, 20 mm depth). Motility was examined on a 37 °C stage of a Nikon E200 microscope under a 10X phase contrast objective (CFI Plan Achro 10X/0.25 Ph1 BM, Nikon). Images were recorded (40 frames at 50 fps) using a CMOS video camera (Basler acA1300-200μm, Basler AG, Ahrensburg, Germany) and analyzed by computer-assisted sperm analysis (CASA, Sperm Class Analyzer version 6.3, Microptic, Barcelona, Spain). Sperm total motility and hyperactivated motility was quantified simultaneously. Over 200 motile sperm were analyzed for each trial.

### RNA isolation

For bulk RNA-sequencing and RT-qPCR, total RNA was extracted from pelleted cells and dissected seminiferous tubules by homogenizing in 1 ml of TRIzol reagent (Invitrogen, 15596026). After 5 mins of incubation, the lysate was mixed with 200μl of chloroform and centrifuged at 12,000 xg for 15 mins at 4°C. The aqueous phase was mixed with an equal volume of 100% ethanol and transferred to an RNeasy MinElute spin column (Qiagen, 74104) for purification according to the manufacturer’s instructions. The quality of the eluted RNA was assessed by 28S/18S ribosomal ratios and RNA integrity number (RIN) using an Agilent Bioanalyzer.

### Real-time quantitative PCR

Reverse transcription of 1 μg of total RNA was performed with oligo(dT) and SuperScript III reverse transcriptase (Thermo, 18080051) in a total volume of 20 μl according to the manufacturer’s instructions. Reaction mixtures were incubated in a thermocycler at 65 °C for 10 mins then 50 °C for 50 mins before stopping the reaction at 85 °C for 5 mins. Real-time quantitative PCR (RT-qPCR) assays were performed in a total reaction volume of 20 μl consisting of 4 μl of cDNA diluted 1:5, 0.4 μl of 10 μM forward/reverse primer mix, 10 μl of Power SYBR Green PCR Master Mix (Applied Biosystems, 4367659) and 5.6 μl nuclease-free water. Reactions for each target gene were performed in duplicate in a 96 well plate loaded into an Applied Biosystems QuantStudio 3 Real-Time PCR system. Standard cycling conditions were used: Hold stage (x1): 50 °C for 2 mins, 95 °C for 10 mins; PCR stage (x40): 95 °C for 15 secs, 60 °C for 1 min. Melt curve stage conditions were: 95 °C for 15 secs, 60°C for 1 min, 95 °C for 15 secs. Relative fold change in transcript abundance was calculated using the delta-delta Ct method by normalizing target gene expression levels to *Actb*. Primer sequences used for RT-qPCR are listed in **Table S1**.

### Bulk RNA-sequencing and analysis

Poly(A)-selected RNA-seq libraries prepared using the KAPA mRNA HyperPrep Kit (Roche # 08098123702) and sequenced to a depth of ∼25 million read pairs per sample. Raw sequence files were filtered and their quality was assessed using FASTX-toolkit and FastQC. Filtered paired-end reads were matched to yield a file of common reads which were then pseudoaligned to the mouse (mm39) transcriptome from Ensembl release 107 (56) and quantified using Kallisto v0.45.0 (57). Resulting transcript counts were summed to gene level and differentially expressed genes were called using DESeq2 (58). Genes with < 10 TPM were filtered from the analysis. Genes with *P* values ≤ 0.05 were considered differentially expressed. A fold change cut-off of ≥1.5 was used to define strongly differentially expressed genes.

### Single cell RNA-sequencing and analysis

Raw count matrices (10X Genomics) for the ‘WT’ and ‘cKO’ mouse were imported to Seurat 4.1.0 (27) and combined into one Seurat object. No explicit integration step was performed because principal component analysis indicated high initial similarity between samples. Reads were filtered to exclude cells likely to be doublets (nCount_RNA < mean + one standard deviation), dead or dying cells (nFeature_RNA > mean – one standard deviation) or high mitochondrial content (present.mt < mean + standard deviation). This filtered subset was log normalized and scaled. A linear dimensional reduction was performed using principal component analysis (PCA), and 20 dimensions was chosen as optimum based on inspection of Scree plots. Clusters were identified at a resolution of 0.5, and the “RunUMAP()” function was used to further investigate the dataset. Three of the initial clusters had a low number of detected molecules per cell, indicating dead or dying cells, and did not have any top genes associated with the transition from spermatogonia to elongating spermatid. Therefore, these three clusters were removed, and the data was normalized, scaled, and dimensionally reduced again using the same parameters described above.

### Chromatin accessibility analysis

Chromatin accessibility was assayed with 100,000 cells per sample. Libraries were prepared using the Active Motif ATAC-Seq kit (53150) according to the manufacturer’s instructions except 11 PCR cycles were used instead of the recommended 10 cycles. Libraries were sequenced to a depth of ∼50 million paired-end 100bp reads per sample. Adapter trimming was performed using Cutadapt (59). Trimmed reads were aligned to the mouse genome (mm10) using Bowtie2 in very sensitive mode (60). Subsequent processing was performed according to the workflow previously described by Reske, Wilson (36). Regions of differential chromatin accessibility were called using *csaw* workflow “method IV” described by Reske *et al*. (2020). Briefly, reads were counted at regions corresponding to the union of peaks identified by default parameters in MACS2 for control and knockdown/knockout samples. Then, low abundance windows were filtered and a non-linear loess-based normalization method was implemented. Resulting count matrices were then subjected to *edgeR* for differential accessibility quantification.

### Functional enrichment analysis

GO enrichment analysis was performed using PANTHER GO annotations (https://doi.org/10.5281/zenodo.4495804 released: February 1, 2021) at the GO Consortium website (Mi et al. 2017; http://www.geneontology.org). Genes considered to be expressed (i.e., >1 TPM) in the relevant sample type were used as the reference list. GSEA was conducted as previously described using version 4.2.3 with default parameters (38, 39).

## Data Availability

Sequencing datasets are deposited at the NCBI Gene Expression Omnibus (GEO) repository under accession number GSE215112.

## Acknowledgments

We thank P. Reddi for the gift of the ACRV1 antibody. We appreciate the technical assistance provided by Aushaq Malla, Delaney Farris, Zachary Smith, and Jake Reske, and the help from the Yale Center for Genome Analysis for high-throughput sequencing. This work was supported by funding from the National Institute of Child Health and Human Development (NICHD, R01HD098128), the Searle Scholars Program, and a Pew Scholar Award to B.J.L. B.W.W. is supported by a postdoctoral fellowship from the Hope Funds for Cancer Research.

## Author contributions

Conceptualization: B.W.W., B.J.L.; Validation: B.W.W.; Formal Analysis: B.W.W., S.R.R., N.D.; Investigation: B.W.W., S.R.R., N.D., X.H., D.G.deR..; Resources: B.J.L.; Writing – Original Draft: B.W.W.; Writing – Review & Editing: B.J.L.; Visualization: B.W.W., S.R.R.; Supervision: B.J.L.; Funding acquisition: B.J.L.

## Supporting Information captions

**S1 Figure. Characterization of scRNA-seq dataset from adult mouse testis and additional in situ hybridization analysis**. (**A**) Graph based clustering of the scRNA-seq data showing the 17 distinct cell populations identified. (**B**) UMAPs of scRNA-seq data (this study) showing the expression for markers of different spermatogenic cell populations and testicular somatic cells. Note that *cKit* expression is shown to highlight the general cell populations retrieved from cKIT-sorting of whole testis (**C**) t-SNE plot showing the expression of *Kdm6a* in mouse testis using scRNA-seq data sourced from Jung et al. 2019. The authors confirmed that t-SNE and UMAP give consistent pseudotime embedding. (**D**) t-SNE plot showing the expression of *Kdm6a* in human testis using scRNA-seq data sourced from Guo et al. 2018. (**E**) Brightfield micrographs showing in situ hybridization for the indicated transcript (pink) in tissue sections of stage XII seminiferous tubules co-stained with hematoxylin. Dashed boxes indicate the regions captured at high magnification below. Scale bar = 50μm. (**F**) UMAPs showing the expression for all lysine demethylases (KDMs) detected in the scRNA-seq dataset.

**S2 Figure. Knockout validation and additional functional assays in *Kdm6a* cKO germ cells**. (**A**) Western blotting of whole lysates from control and *Kdm6a* cKO testes for KDM6A and histone H3 (loading control). (**B**) RT-qPCR for *Kdm7b* and the KDM6A homologs *Uty* and *Kdm6b* in cDNA samples of control and *Kdm6a* cKO testes. (**C**) Quantification of preleptotene cells by visual examination of testes sections stained with hematoxylin and eosin. (**D**) Quantification of sperm motility parameters by computer-assisted sperm analysis (CASA) in control and *Kdm6a* cKO samples under capacitated and uncapacitated conditions. ns = not significant.

**S3 Figure. Quality Assessment of ATAC-seq datasets**. (**A**) Western blotting cytoplasmic and nuclear fractions extracted from wildtype testis for KDM6A, histone H3 (nuclear marker), DDX4 (cytoplasmic germ cell marker), and tubulin (cytoplasmic marker). (**B**) Signal enrichment for regions with ATAC-seq peaks at different genomic features. (**C**) Gene feature distribution of ATAC-enriched regions in control cKIT+ testis cells. (**D**) Genome browser tracks showing representative ATAC-seq peaks detected in cKIT+ testis cells at *Gapdh* (housekeeping gene) and neighboring genes. Numbers to the right represent the set scale range for each sample. (**E**) Gene feature distribution for regions with ATAC-seq peaks in shScr-GC1-SPGs. (**F**) Violin plot showing the expression levels (transcripts per million, TPM) for genes with and without ATAC-seq peaks at the promoter in shScr-GC1-SPGs. (**G**) Genome browser tracks showing representative ATAC-seq peaks at *Gapdh* and neighboring genes in GC1-SPGs. (**H**) Intersection of genome coordinates for ATAC-enriched regions in shScr-GC1-SPGs and control cKIT+ cells.

**S4 Figure. Validation of Transcriptome analysis in *Kdm6a* cKO testis and GC1 cell cultures**. (**A**) Heatmap showing the expression of *Kdm6a* cKO testis scDEGs across control testis cells for each cluster. (**B**) Gene set enrichment analyses of bulk-RNA-seq data from *Kdm6a* cKO testes with differentially expressed genes identified by scRNA-seq (scDEGs). (**C**) Volcano plot showing the changes in gene expression detected in sh*Kdm6a*#1-GC1s. (**D**) Volcano plot showing the changes in gene expression detected in sh*Kdm6a*#2-GC1s. (**E**) Overlap of downregulated (above) and upregulated (below) DEGs identified for GC1-SPGs expressing sh*Kdm6a*#1 or sh*Kdm6a*#2. (**F**) Gene set enrichment analyses of sh*Kdm6a*#1-GC1s expression data with genes associated with differentially accessible regions of chromatin identified by ATAC-seq. Normalized enrichment score (NES). (**G**) List of DEGs detected in sh*Kdm6a*-GC1-SPGs that are shared with scDEGs from *Kdm6a* cKO testis.

**S5 Figure. Validation of *Kdm6a* expression status and additional gene set enrichment analysis in *Kdm6a* F1 and *Kdm6a* F2 testes**. (**A**) RT-qPCR analysis for *Kdm6a* expression normalized to *Actb* in whole testis samples from control, *Kdm6a* cKO, *Kdm6a* F1, and *Kdm6a* F2 mice (n =3). Errors bars = standard error of the mean. (**B**) Normalized enrichment scores from gene set enrichment analysis (GSEA) of *Kdm6a* F1 testis expression data with upregulated (red) and downregulated (blue) differentially expressed genes (scDEGs) identified for different testis cell populations in *Kdm6a* cKO mice. * FDR q-value = < 0.05. (**C**) GSEA plots showing the enrichment of *Kdm6a* F1 and *Kdm6a* F2 testis expression data for *Kdm6a* cKO scDEGs identified for cluster 4 and cluster 5. Normalized enrichment score = NES.

**S1 Table**. Primer sequences used in this study.

**S2 Table**. Antibodies used in this study.

**S1 Dataset**. ATAC-seq peaks.

**S2 Dataset**. scRNA-seq DEGs by cluster.

**S3 Dataset**. GC1 DEGs.

**S4 Dataset**. Testis DEGs in the F1 and F2 generations.

