## Supplementary material for "KDM6A/UTX promotes spermatogenic gene expression across generations but is dispensable for male fertility": S1 Figure

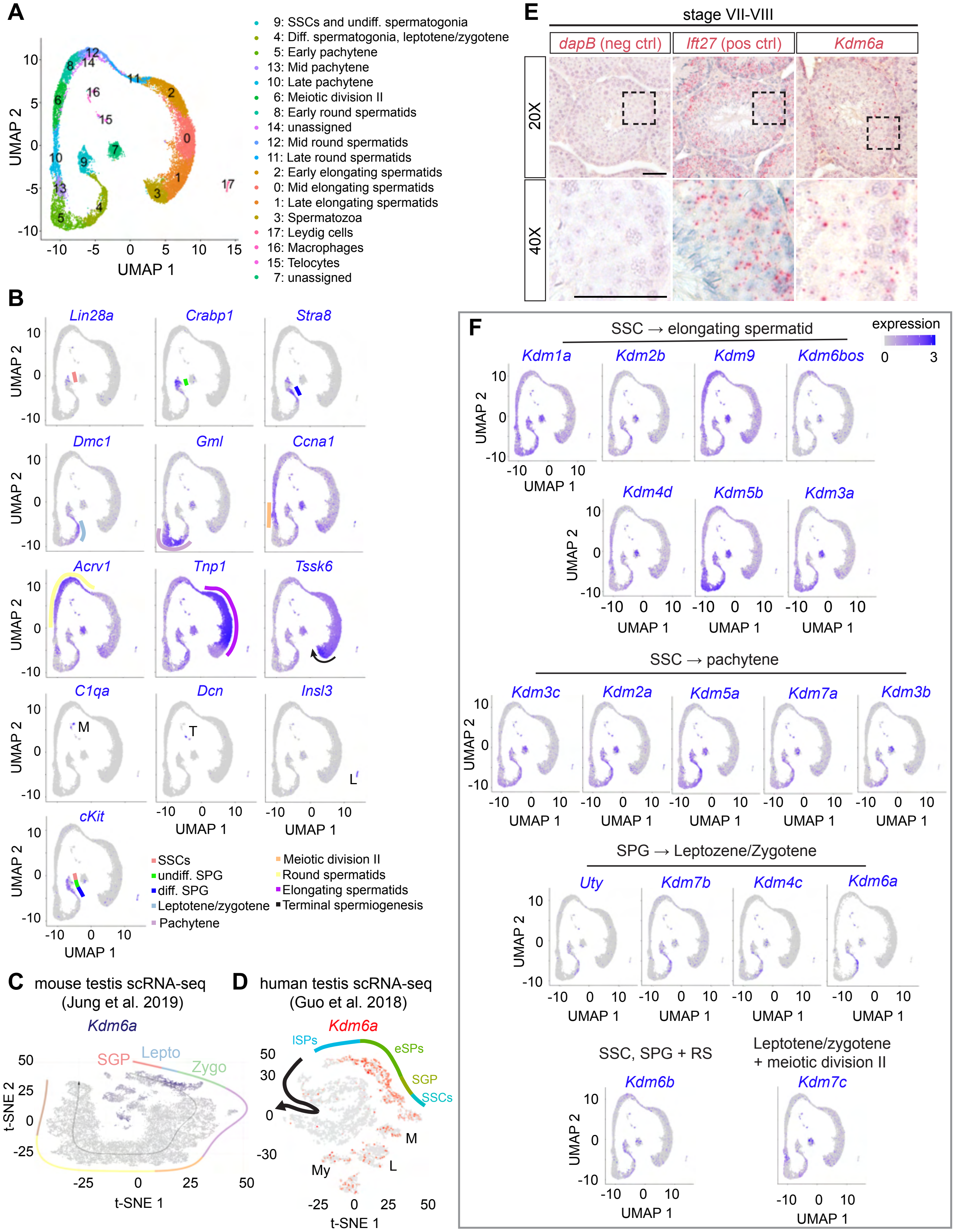


**Figure S1. Characterization of scRNA-seq dataset from adult mouse testis and additional in situ hybridization analysis.** (**A**) Graph based clustering of the scRNA-seq data showing the 17 distinct cell populations identified. (**B**) UMAPs of scRNA-seq data (this study) showing the expression for markers of different spermatogenic cell populations and testicular somatic cells. Note that *cKit* expression is shown to highlight the general cell populations retrieved from cKIT+ sorting of whole testis (**C**) t-SNE plot showing the expression of *Kdm6a* in mouse testis using scRNA-seq data sourced from Jung et al. 2019. The authors confirmed that t-SNE and UMAP give consistent pseudotime embedding. (**D**) t-SNE plot showing the expression of *Kdm6a* in human testis using scRNA-seq data sourced from Guo et al. 2018. (**E**) Brightfield micrographs showing in situ hybridization for the indicated transcript (pink) in tissue sections of stage XII seminiferous tubules co-stained with hematoxylin. Dashed boxes indicate the regions captured at high magnification below. Scale bar = 50μm. (**F**) UMAPs showing the expression for all lysine demethylases (KDMs) detected in the scRNA-seq dataset.
