## Supplementary material for "KDM6A/UTX promotes spermatogenic gene expression across generations but is dispensable for male fertility": S2 Figure

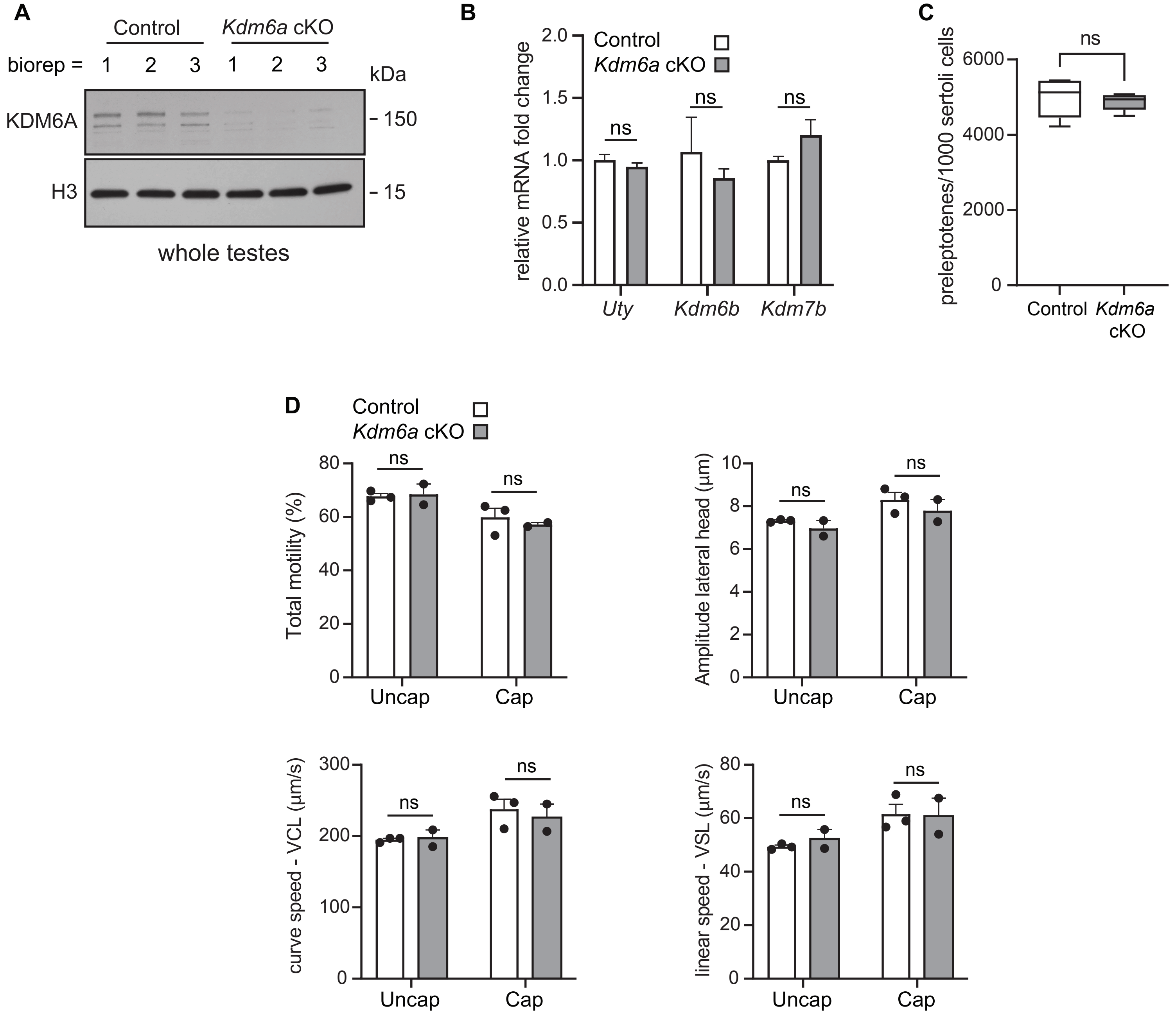


**Figure S2. Knockout validation and additional functional assays in *Kdm6a* cKO germ cells.** (**A**) Western blotting of whole lysates from control and *Kdm6a* cKO testes for KDM6A and histone H3 (loading control). (**B**) RT-qPCR for *Kdm7b* and the KDM6A homologs *Uty* and *Kdm6b* in cDNA samples of control and *Kdm6a* cKO testes. (**C**) Quantification of preleptotene cells by visual examination of testes sections stained with hematoxylin and eosin. (**D**) Quantification of sperm motility parameters by computer-assisted sperm analysis (CASA) in control and *Kdm6a* cKO samples under capacitated and uncapacitated conditions. ns = not significant.
