## Supplementary material for "KDM6A/UTX promotes spermatogenic gene expression across generations but is dispensable for male fertility": S3 Figure

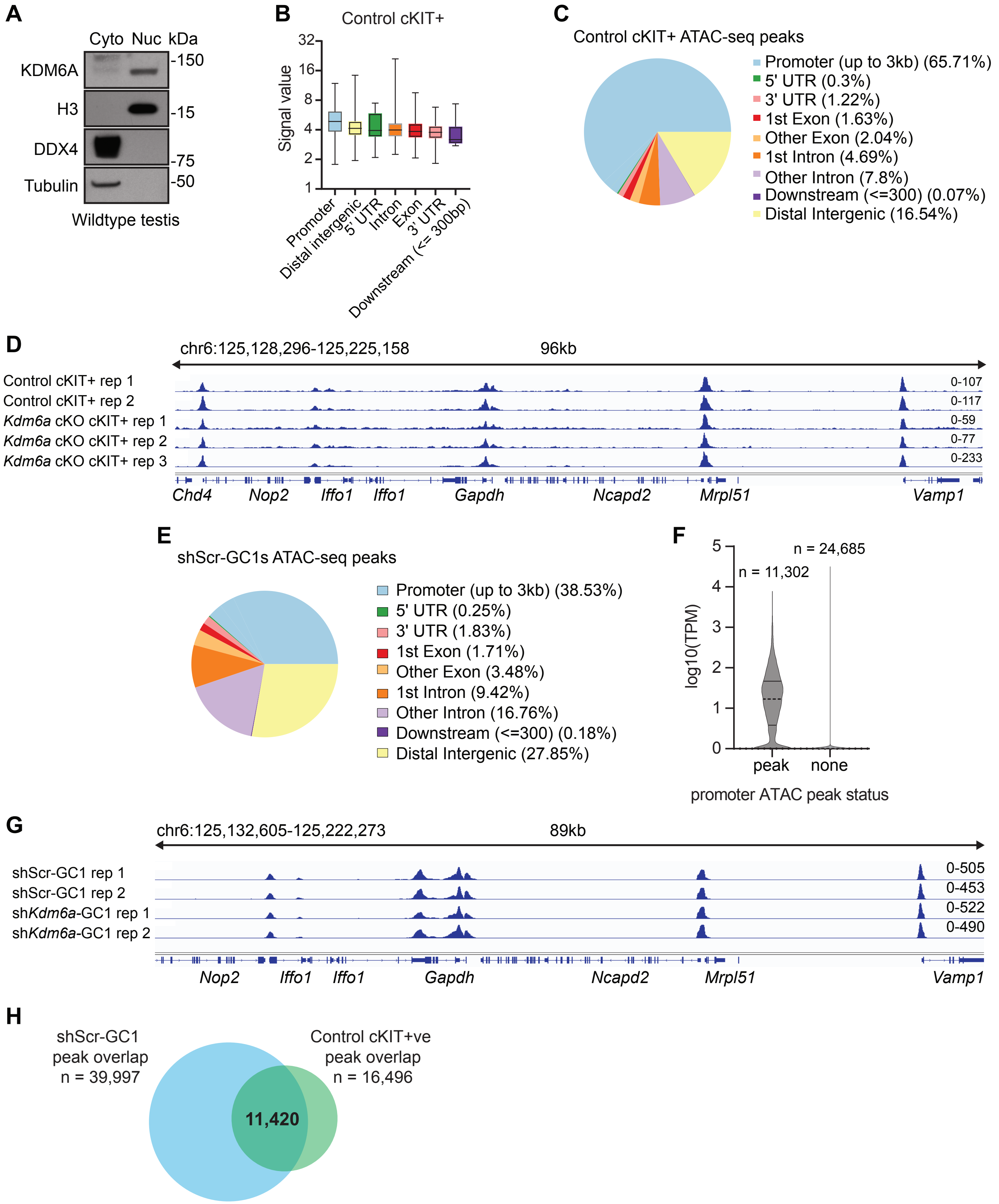


**Figure S3. Quality assessment of ATAC-seq datasets.** (**A**) Western blotting cytoplasmic and nuclear fractions extracted from wildtype testis for KDM6A, histone H3 (nuclear marker), DDX4 (cytoplasmic germ cell marker), and tubulin (cytoplasmic marker). (**B**) Signal enrichment for regions with ATAC-seq peaks at different genomic features. (**C**) Gene feature distribution of ATAC-enriched regions in control cKIT+ testis cells. (**D**) Genome browser tracks showing representative ATAC-seq peaks detected in cKIT+ testis cells at *Gapdh* (housekeeping gene) and neighboring genes. Numbers to the right represent the set scale range for each sample. (**E**) Gene feature distribution for regions with ATAC-seq peaks in shScr-GC1-SPGs. (**F**) Violin plot showing the expression levels (transcripts per million, TPM) for genes with and without ATAC-seq peaks at the promoter in shScr-GC1-SPGs. (**G**) Genome browser tracks showing representative ATAC-seq peaks at *Gapdh* and neighboring genes in GC1-SPGs. (**H**) Intersection of genome coordinates for ATAC-enriched regions in shScr-GC1-SPGs and control cKIT+ cells.
