## Supplementary material for "KDM6A/UTX promotes spermatogenic gene expression across generations but is dispensable for male fertility": S4 Figure

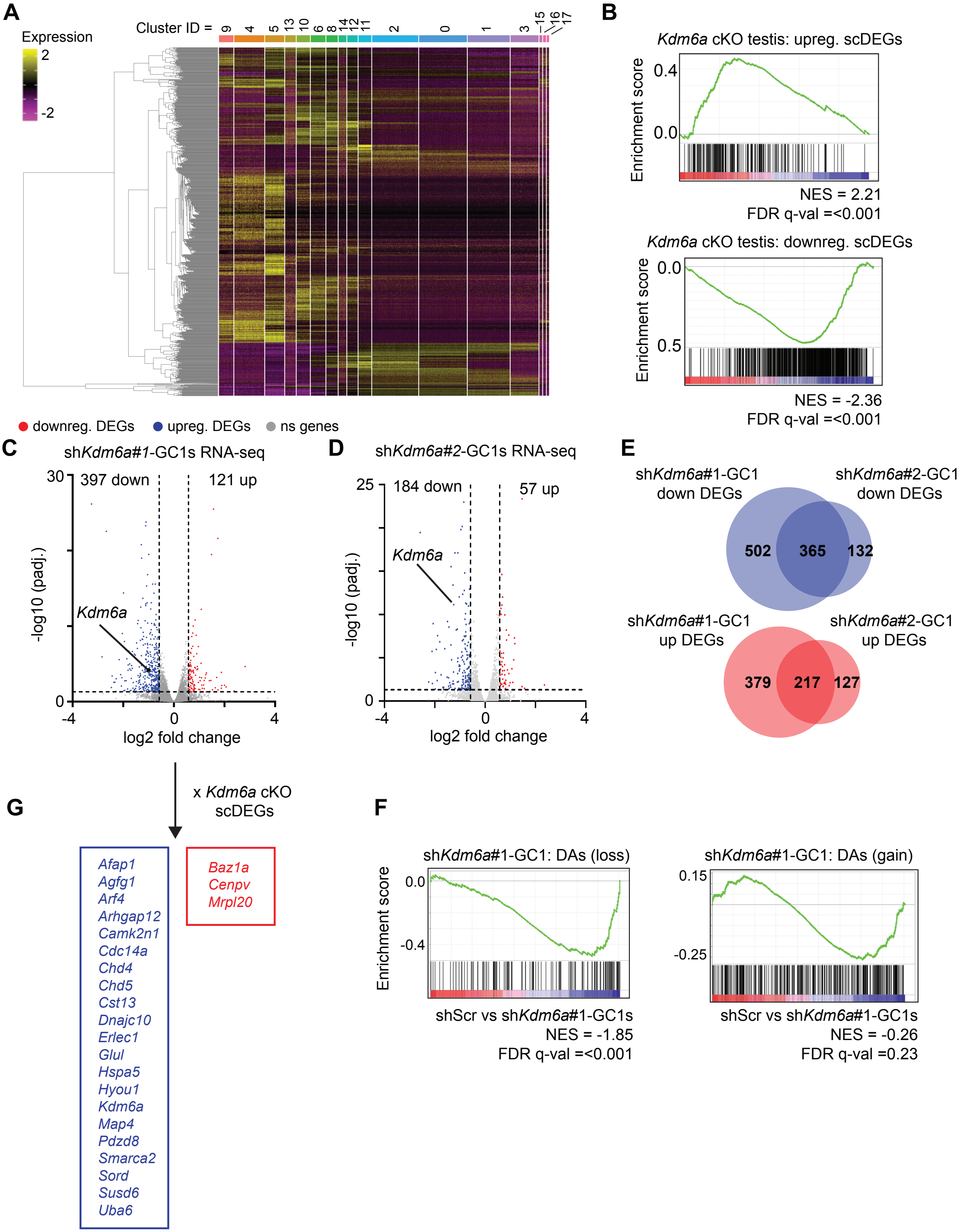


**Figure S4. Validation of transcriptome analysis in Kdm6a cKO testis and GC1 cell cultures.** (**A**) Heatmap showing the expression of *Kdm6a* cKO testis scDEGs across control testis cells for each cluster. (**B**) Gene set enrichment analyses of bulk-RNA-seq data from *Kdm6a* cKO testes with differentially expressed genes identified by scRNA-seq (scDEGs). (**C**) Volcano plot showing the changes in gene expression detected in *shKdm6a*#1-GC1s. (**D**) Volcano plot showing the changes in gene expression detected in *shKdm6a*#2-GC1s. (**E**) Overlap of downregulated (above) and upregulated (below) DEGs identified for GC1-SPGs expressing *shKdm6a*#1 or *shKdm6a*#2. (**F**) Gene set enrichment analyses of *shKdm6a*#1-GC1s expression data with genes associated with differentially accessible regions of chromatin identified by ATAC-seq. Normalized enrichment score (NES). (**G**) List of DEGs detected in *shKdm6a*-GC1-SPGs that are shared with scDEGs from *Kdm6a* cKO testis.
