## Supplementary material for "KDM6A/UTX promotes spermatogenic gene expression across generations but is dispensable for male fertility": S5 Figure

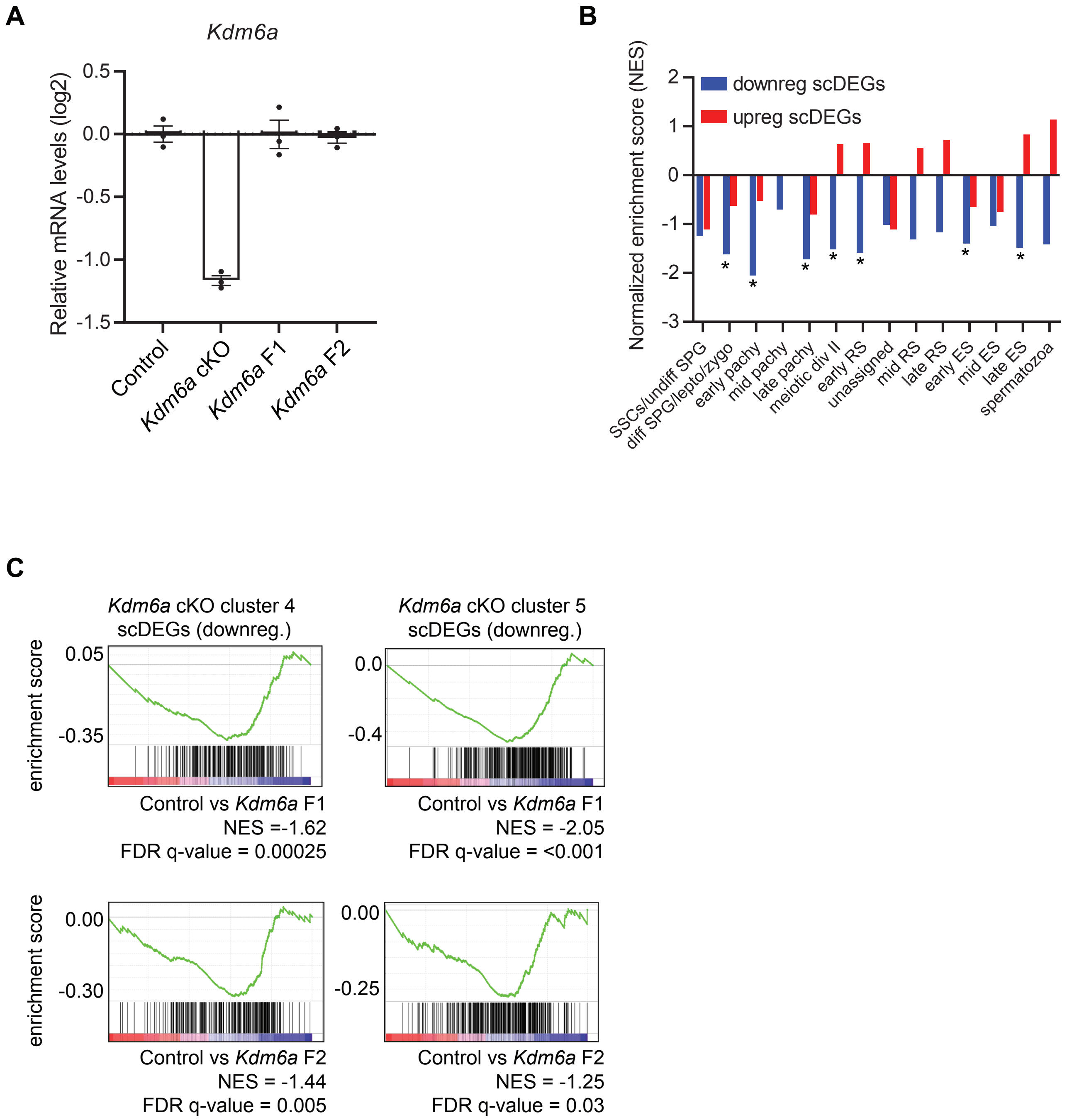


**Figure S5. Validation of *Kdm6a* expression status and additional gene set enrichment analysis in *Kdm6a* F1 and *Kdm6a* F2 testes.** (**A**) RT-qPCR analysis for *Kdm6a* expression normalized to *Actb* in whole testis samples from control, *Kdm6a* cKO, *Kdm6a* F1, and *Kdm6a* F2 mice (n =3). Errors bars = standard error of the mean. (**B**) Normalized enrichment scores from gene set enrichment analysis (GSEA) of *Kdm6a* F1 testis expression data with upregulated (red) and downregulated (blue) differentially expressed genes (scDEGs) identified for different testis cell populations in *Kdm6a* cKO mice. * FDR q-value = < 0.05. (**C**) GSEA plots showing the enrichment of *Kdm6a* F1 and *Kdm6a* F2 testis expression data for *Kdm6a* cKO scDEGs identified for cluster 4 and cluster 5. Normalized enrichment score = NES.
